# Crustaceans as hosts of parasites throughout the Phanerozoic

**DOI:** 10.1101/505495

**Authors:** Adiёl A. Klompmaker, Cristina M. Robins, Roger W. Portell, Antonio De Angeli

## Abstract

The fossil record of crustaceans as hosts of parasites has yielded three confirmed associations: epicaridean isopod-induced swellings on Jurassic–Recent decapod crustaceans, feminization of Cretaceous and Miocene male crabs possibly caused by rhizocephalan barnacles, and presumed pentastomids on/in Silurian ostracods. Cestode platyhelminth hooks and swellings by entoniscid isopods may be recognized in the future. Relative to 2014, we report an increase of 41% to 124 fossil decapod species with epicaridean-induced swellings in the branchial chamber (ichnotaxon *Kanthyloma crusta*). Furthermore, using a Late Jurassic (Tithonian) decapod assemblage from Austria, we find (1) no correlation between genus abundance and prevalence of *K. crusta*, (2) host preference for some galatheoid taxa (as for a mid-Cretaceous assemblage from Spain), and (3) a larger median size of parasitized versus non-parasitized specimens for two selected species. The latter result may be caused by infestation throughout ontogeny rather than exclusively in juveniles and/or possible selection for the larger sex.

## 1. Introduction

Extant crustaceans (or pancrustaceans) act as both parasites and hosts of parasites (e.g., Yamaguti 1963; Cressey 1983; Boxshall et al. 2005; Boyko and Williams 2011; Trilles and Hipeau-Jacquotte 2012; Smit et al. 2014; Klompmaker and Boxshall 2015; Boxshall and Hayes in press). For this paper, we define parasitism as a symbiotic relationship in which the parasite is nutritionally dependent on the host for at least part of its life cycle and has a negative impact on the fitness of the host (cf. Combes 2001; Tapanila 2008). Over 7000 extant crustaceans are intermediate and final hosts to parasites from diverse clades such as acanthocephalans, cestodes, crustaceans (e.g., copepods, cirripedes, isopods, Tantulocarida), digenean trematodes, monogeneans, nematodes, and protists (e.g., Boxshall et al. 2005; Klompmaker and Boxshall 2015; Boxshall and Hayes in press). Representatives of nearly all to all major clades of crustaceans serve as hosts today, including amphipods, branchiopods, cirripeds, copepods, decapods, euphausiaceans, mysidaceans, ostracods, peracarids, and stomatopods (e.g., Boxshall et al. 2005; Klompmaker and Boxshall 2015; Boxshall and Hayes in press). The impact of parasitism on crustaceans is likely to be enormous, but understudied in the context of whole ecosystems.

The fossil record of parasitism involving crustaceans has been reviewed recently (Klompmaker and Boxshall 2015, for crustaceans as parasites and hosts; Haug et al. this volume, for crustaceans as parasites). Three instances of parasites in crustacean hosts have been reported thus far (Fig. 1). This low number and the fact that the stratigraphic coverage of two of these records is spotty can be ascribed to a combination of the small size of parasites, the low preservation potential of parasites due to their general lack of a hard skeleton, not all parasites leave recognizable traces, and the lack of targeted research. One notable exception is epicaridean isopods, which cause characteristic swellings on decapod crustacean carapaces. This association represents a nearly continuous record since the Jurassic (Klompmaker et al. 2014; Klompmaker and Boxshall 2015), presenting an ideal model system to study various aspects of parasitism through time.

**Figure 1.**
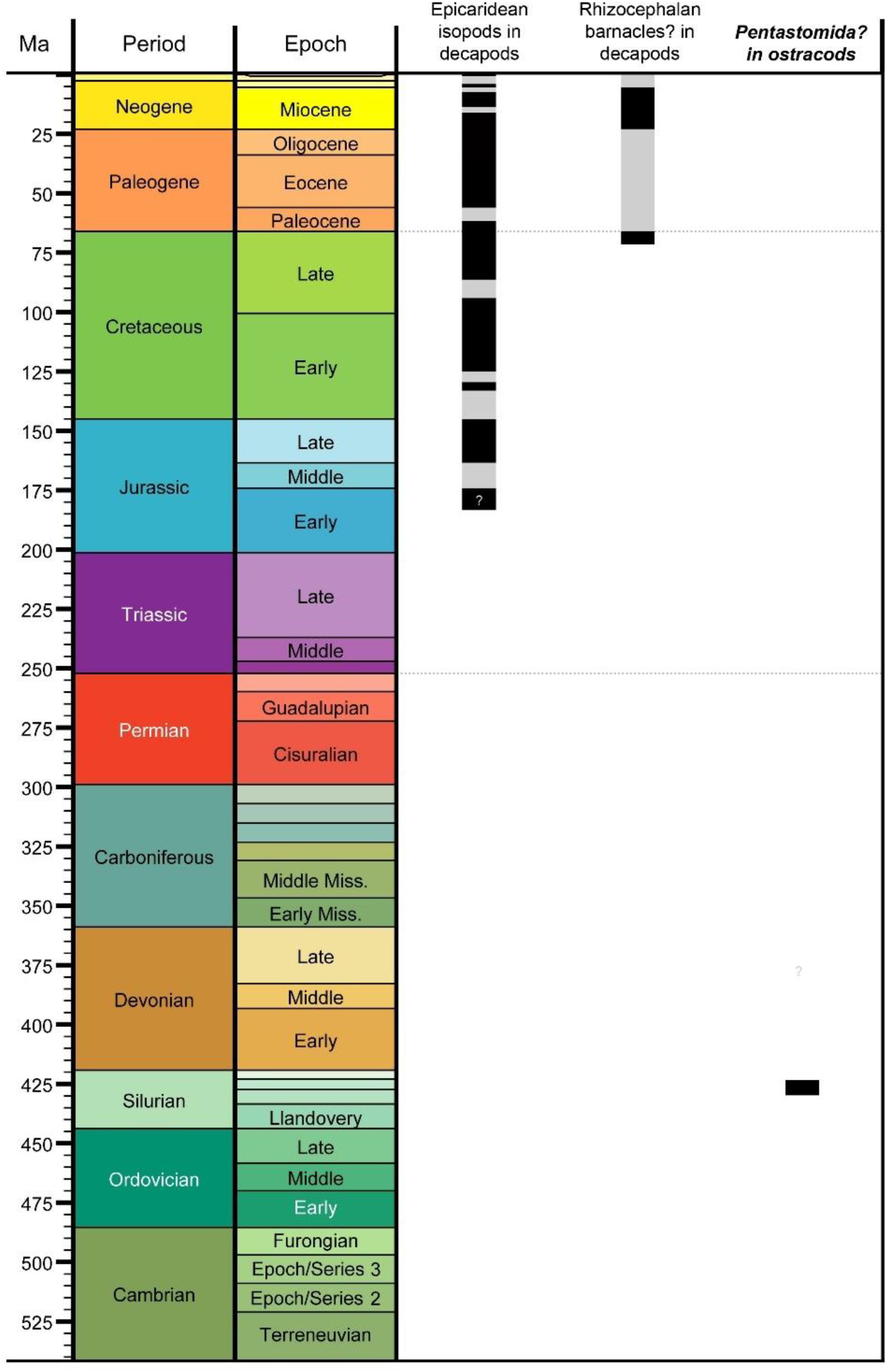
Stratigraphic ranges of crustaceans as hosts of parasites. Black parts represent known occurrences, while grey parts are inferred occurrences. Black occurrences have been plotted at the stage-level where possible. The category in bold italic font is based on body fossils of the parasite; others are based on morphologies inferred to have been caused by parasites in the host taxon. Timescale on left produced with TSCreator 6.3 (http://www.tscreator.org). Modified and updated from Klompmaker and Boxshall (2015: fig. 12).

The goal of this paper is to re-review the fossil record of crustaceans as hosts of parasites because ample new evidence has been found in recent years. We focus primarily on evidence of epicaridean isopod parasites in decapod crustaceans by (1) presenting a substantially expanded list of infested decapod species and (2) using a vast Late Jurassic assemblage to test the relationship between taxon abundance and infestation percentage, assess host preference, and evaluate the size of parasitized versus non-parasitized specimens for two species. Subsequently, we briefly review the claimed evidence for parasitism of (1) rhizocephalan barnacles in fossil decapod crustaceans, (2) ciliates on ostracods, and (3) pentastomids on ostracods. Finally, modern parasitism on crustaceans with preservation potential are discussed.

Institutional abbreviations: MAB = Oertijdmuseum, Boxtel, The Netherlands; MCV = Museo Civico D. Dal Lago, Valdagno, Vicenza, Italy; NHMW = Naturhistorisches Museum Wien, Vienna, Austria; UF =Florida Museum of Natural History at the University of Florida, Gainesville, Florida, USA.

## 2. Isopod swellings in decapod crustaceans

### 2.1. General information

Epicaridean isopods (Bopyroidea and Cryptoniscoidea) parasitize calanoid copepods as intermediate hosts and other crustaceans as final hosts, following two larval stages (e.g., Williams and Boyko 2012). Approximately 800 species are known and they are found today in all oceans (e.g., Williams and Boyko 2012). The phylogenetic relationship of epicaridean families has recently been clarified using 18S rDNA, resulting in five accepted families to date (Boyko et al. 2013). Unlike the endoparasitic Cryptoniscoidea (Dajidae and Entophilidae), Bopyroidea (Ionidae, Bopyridae, and Entoniscidae) are ectoparasitic and their female individuals often create swellings in the cuticle of one or rarely both branchial chambers of decapod crustaceans (e.g., Markham 1986; Boyko and Williams 2009; Williams and Boyko 2012). Epicaridean females feed on hemolymph and ovarian fluids after piercing the inner cuticle of the host (e.g., Bursey 1978; Lester 2005), while the much smaller, less deformed, less modified males attach themselves to females subsequently for reproduction and do not form swellings in the cuticle. Effects on the host are manifold and can be dramatic (Table 1).

**Table 1.**
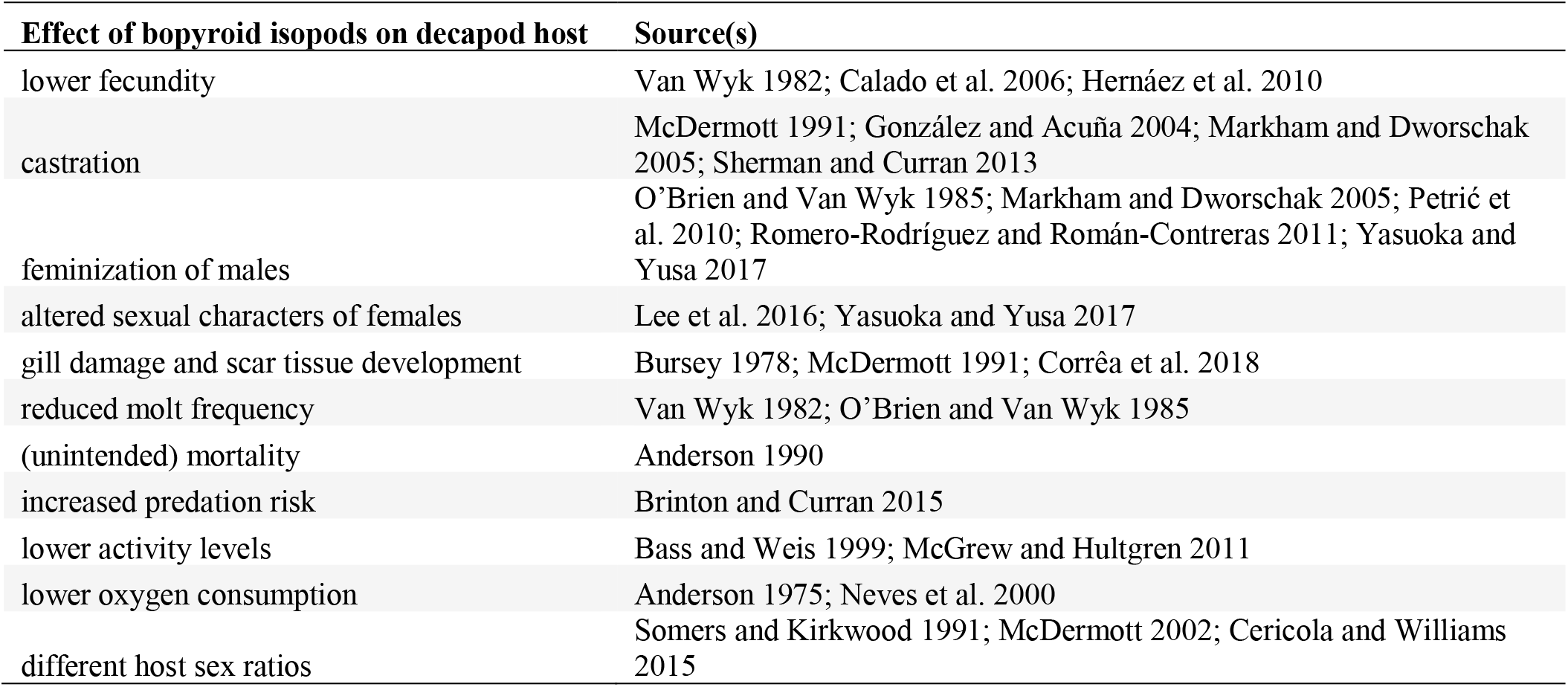
Parasitic epicarideans cause a variety of negative effects on their decapod hosts. Sources are not comprehensive.

The body fossil record of epicarideans is nearly non-existent. No adults have been found in decapod body chambers thus far due to their low preservation potential, as also shown experimentally (Klompmaker et al. 2017; Fig. 2). Only some epicaridean larvae have been found in Miocene and Late Cretaceous amber (Serrano-Sánchez et al. 2016; Néraudeau et al. 2017; Schädel et al. 2018). However, the swellings in the oft-calcified cuticle of decapod hosts can be preserved and detected relatively easily in the fossil record (e.g., Wienberg Rasmussen et al. 2008; Ceccon and De Angeli 2013; Robins et al. 2013; Klompmaker et al. 2014; Klompmaker and Boxshall 2015; Hyžný et al. 2015). These swellings are referred to the ichnotaxon *Kanthyloma crusta* Klompmaker et al., 2014 (see also Klompmaker and Boxshall 2015). The earliest known swelling is in a lobster from the Early Jurassic (Toarcian) of Indonesia, but this record is somewhat doubtful (Wienberg Rasmussen et al. 2008; Klompmaker et al. 2014). The Late Jurassic (Oxfordian) yields the first definite examples, suggesting coevolution between epicarideans and their decapod host for at least 160 million years. Most infested fossil species are brachyurans and galatheoids, taxa that today are parasitized primarily by Ionidae and Bopyridae, respectively (Markham 1986; Boyko and Williams 2009).

**Figure 2.**
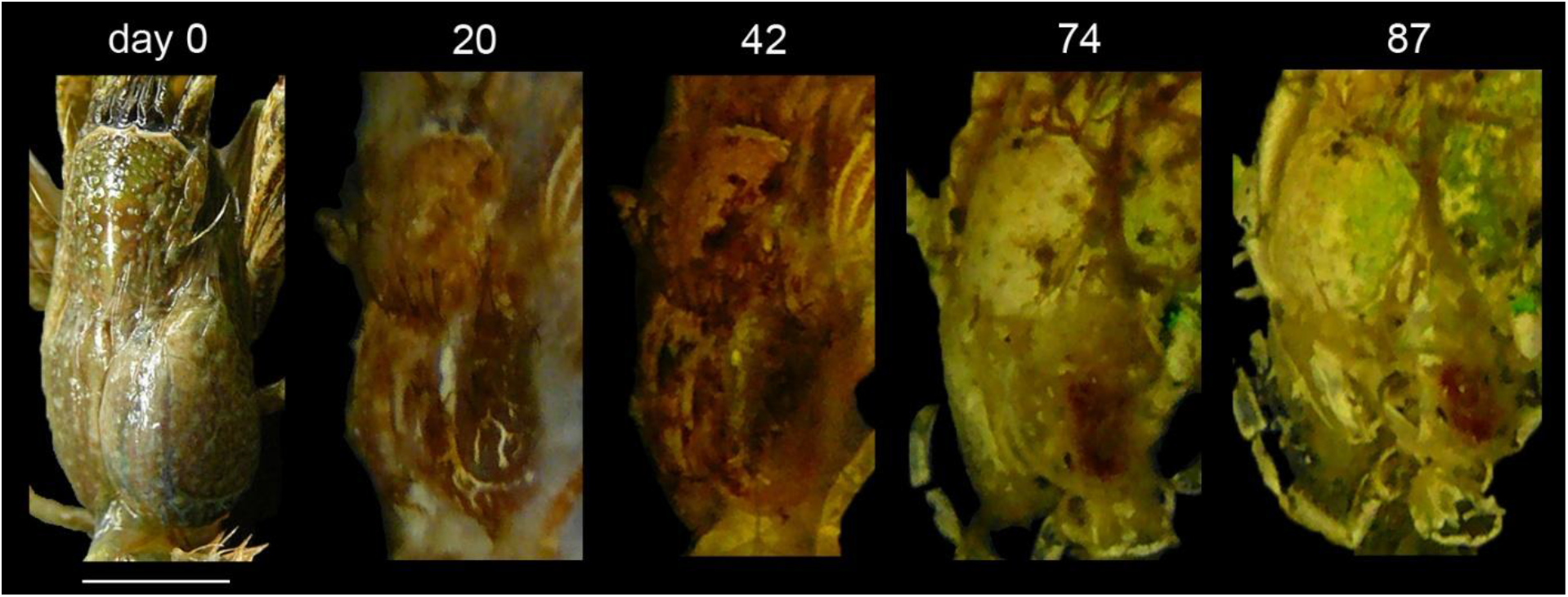
Decay of the carapace of the modern hermit crab *Clibanarius vittatus* (Bosc, 1802) and its parasitic bopyrid [most likely *Bopyrissa wolffi* Markham, 1978, see Markham (1978) and McDermott et al. (2010)] in the swollen right branchial chamber. The carapace is tilted to the left to better observe the decay of the parasite. The bopyrid body outline is still visible on day 20, but much less so subsequently when a smaller red spot is observed. For experimental setup and conditions see Klompmaker et al. (2017). 1^st^, 2^nd^, and 4^th^ image modified from same reference. Scale bar: 5.0 mm wide.

### 2.2. Global Meso- and Cenozoic data

Fossil decapod crustacean species with at least one specimen containing a swelling in the branchial region attributed to an epicaridean parasite have been listed previously (Van Straelen 1928: p. 51; Van Straelen 1931: p. 56; Houša 1963: p. 110; Förster 1969: p. 53–54; Markham 1986: table 2; Wienberg Rasmussen et al. 2008: table 2; Klompmaker et al. 2014: table 3). The most recent report identified 88 parasitized species. Targeted research on Meso-Cenozoic decapod assemblages for evidence of *Kanthyloma crusta* since the summer of 2017, published research since 2014, and new discoveries of infested species in older literature has led to an increase of 41% of known infested fossil species, totaling of 124 infested species (Fig. 3, for examples of new and some known occurrences). This remarkable rise and the total number of infested decapod species today exceeding 550 (Boyko and Williams 2009: fig. 6, and subsequent papers) suggest that more infested fossil taxa are to be expected.

**Figure 3.**
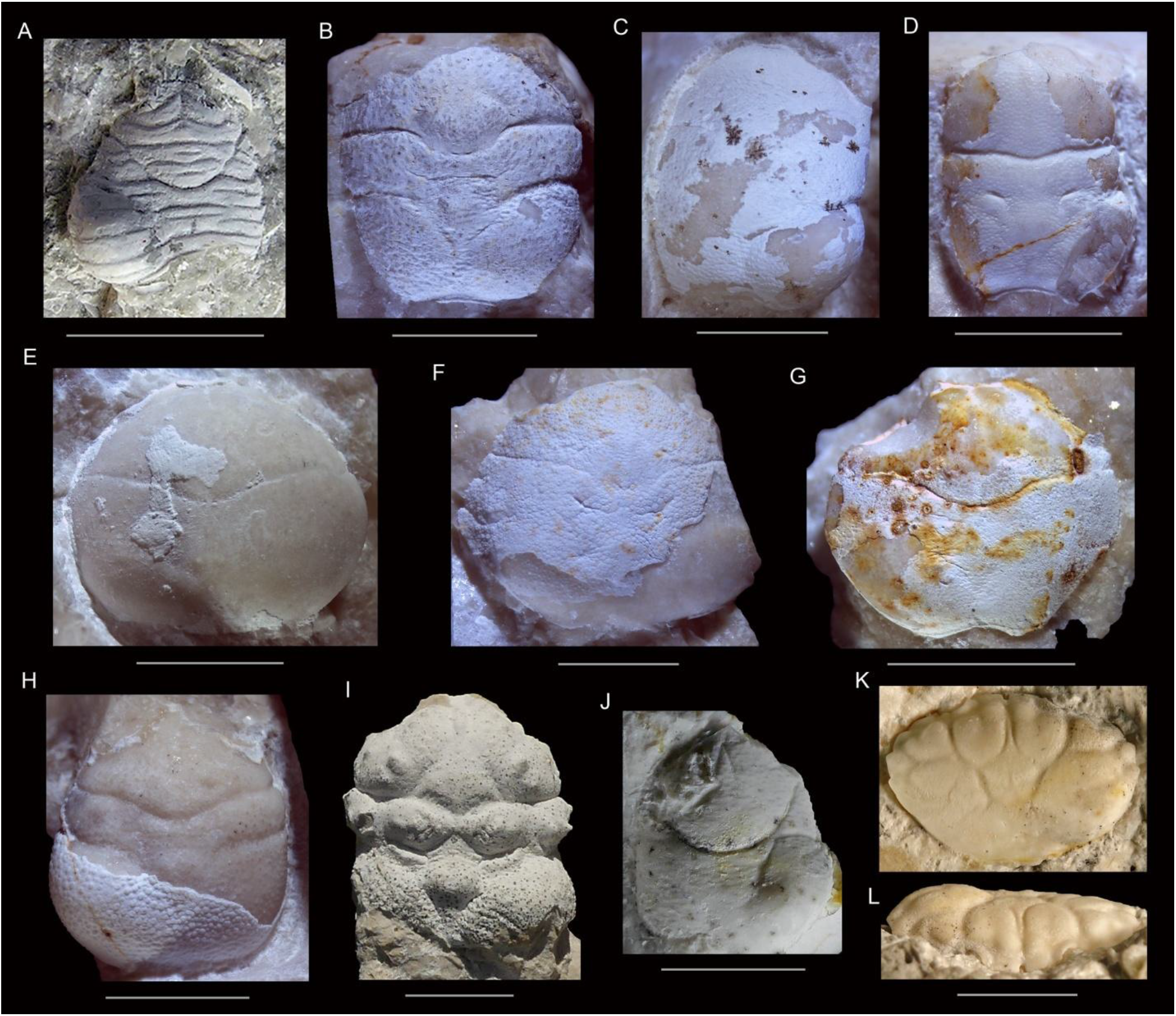
Fossil decapod crustacean carapaces with isopod-induced swellings (ichnotaxon *Kanthyloma crusta*) in one of the branchial chambers. A. *Galathea valmaranensis* De Angeli and Garassino, 2002, from the Oligocene (Rupelian) of Sant’Urbano, Italy (left side), MCV.17/0049. B. *Pithonoton marginatum* von Meyer, 1842 [sensu Wehner (1988: pl. 6.1)] (right side), NHMW 1990/0041/1373. C. *Pithonoton* cf. *P. rusticum* Patrulius, 1966 (right side), NHMW 2017/0089/0028. D. *Longodromites angustus* (Reuss, 1858) (right side), NHMW 1990/0041/2600. E. *Cyclothyreus cardiacus* Schweitzer and Feldmann, 2009a (left side), NHMW 2017/0089/0026. F. *Distefania oxythyreiformis* (Gemmellaro, 1869) (right side), NHMW 2017/0089/0002. G. *Cycloprosopon* cf. *C. octonarium* Schweitzer and Feldmann, 2010 (left side), NHMW 1990/0041/1455. H. *Coelopus hoheneggeri* (Moericke, 1889) (left side), NHMW 1990/0041/1519. I. *Abyssophthalmus spinosus* (von Meyer, 1842) from the Late Jurassic (Kimmeridgian) of the Plettenberg, Germany (right side), MAB k3612. J. *Galatheites* cf. *G. diasema* Robins et al., 2016 (right side), NHMW 1990/0041/0163. K, L. *Panopeus nanus* Portell and Collins, 2004, from the early Miocene of Jamaica (right side), UF 288470. C–H, J from the Late Jurassic (Tithonian) near Ernstbrunn, eastern Austria. All dorsal views, except for L (frontal view). Scale bars: 5.0 mm wide.

Most infested fossil species represent Brachyura and Galatheoidea, and are primarily found in Europe (100/123 or 81%, Fig. 4B). The highest number of infested species on the epoch-level is known from the Late Jurassic (Fig. 4A). A Late Jurassic peak was also shown for the percentage of infested decapod species per epoch in the Mesozoic (Klompmaker et al. 2014: fig. 6). Although they argued that elevated collecting and reporting of *Kanthyloma* in the Late Jurassic may explain part of the peak, they found a biological explanation more likely. Variable reporting may indeed be a factor because there are multiple papers specifically dedicated to this type of parasitism in the Late Jurassic (Remeš 1921; Bachmayer 1948; Houša 1963; Radwański 1972), while only two papers have focused on *Kanthyloma* from younger, pre-Holocene epochs (Ceccon and De Angeli 2013: Eocene; Klompmaker et al. 2014: mid-Cretaceous). Whether this type of parasitism is truly reaching its peak in the Late Jurassic can only be assessed by studying the infestation prevalence on the finest possible scale using specimens from many assemblages across time and space. This study is currently ongoing.

**Figure 4.**
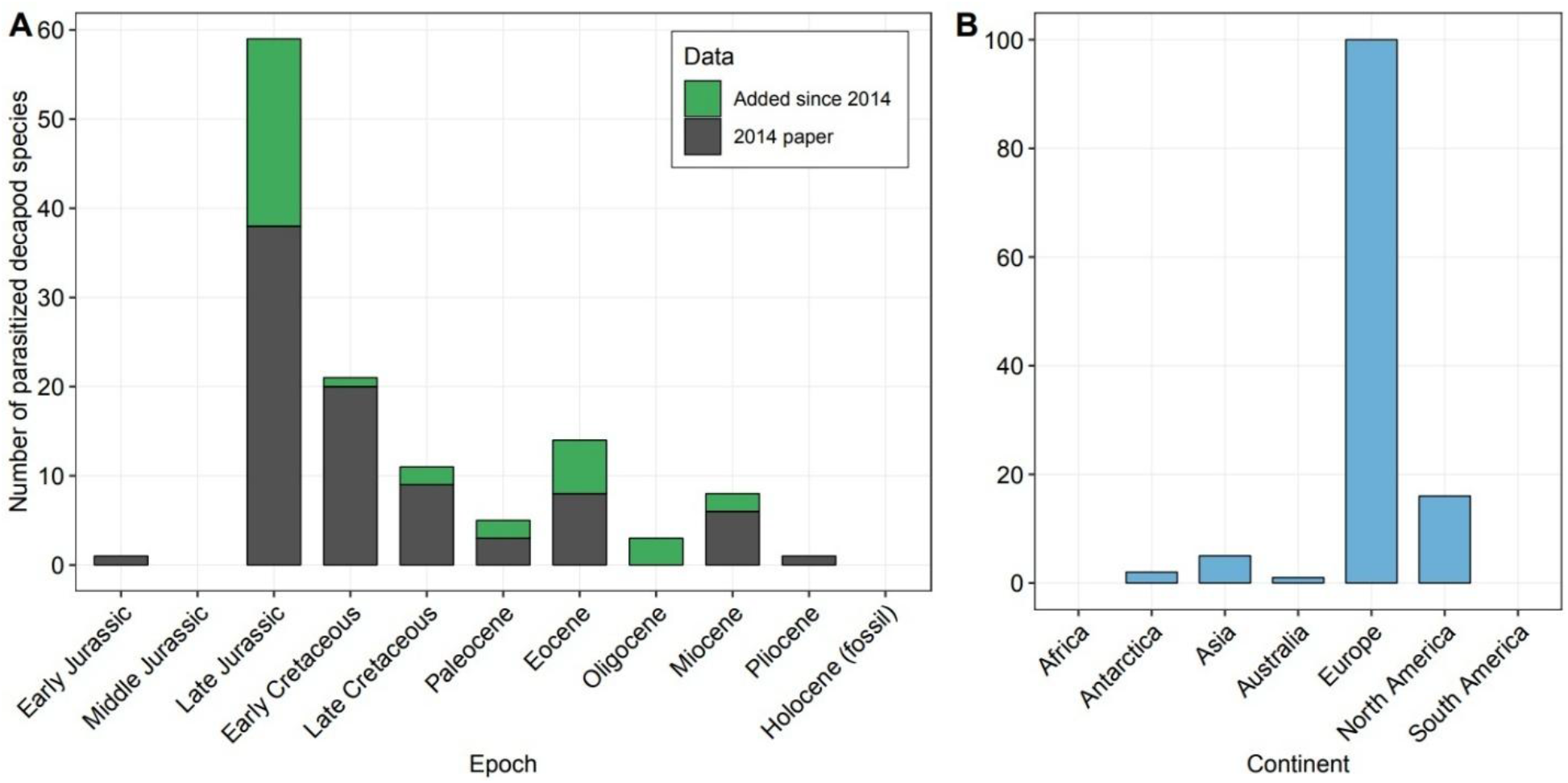
The number of decapod crustacean species parasitized by epicaridean isopods. A. Raw epoch-level data split before and after Klompmaker et al. (2014: table 3), total n = 124. Targeted research has led to 41% more decapod species that were infested. B. Data split by continent.

### 2.3. Abundance vs infestation percentage per taxon

Multiple studies have shown that host density positively affects the transmission rate of parasites. For example, there is a positive correlation between mammal host density across many species and strongylid nematode parasite abundance in modern ecosystems (Arneberg et al. 1998). For decapods, host abundance was the best predictor of bopyrid infestation prevalence in modern carid shrimp from Florida (Briggs et al. 2017). An Early Cretaceous decapod assemblage from Koskobilo in northern Spain yielded a significant correlation between taxon abundance and epicaridean infestation percentage on the species- and genus-levels (Klompmaker et al. 2014: fig. 4A, 4B). These fossil specimens were collected predominantly from the southern wall of the Koskobilo quarry, minimizing the influence of possible spatial and temporal variation. However, they called for more research because of the limited sample size of specimens of many taxa and the fact that the correlation appears driven by one taxon.

To address these issues, the latest Jurassic – earliest Cretaceous Ernstbrunn coral-associated assemblage (also called the Bachmayer Collection) from eastern Austria (e.g., Schneider et al. 2013) was used. This decapod collection consists of ~6900 specimens according to the latest counts (pers. comm. A. Kroh to AAK, November 2018). Bachmayer (1945) listed four localities in the Ernstbrunn Limestone in which decapods were found (Dörfles I, Dörfles Werk II, Klafterbrunn I, and Klement I), but decapods were only abundant in Dörfles I and rare in the other three localities. The Dörfles exposures are considered to be middle to late Tithonian in age based on ammonite stratigraphy (Zeiss 2001; Schweitzer and Feldmann 2009b). All specimens of this collection were identified to the genus- and family-levels, where possible. Species-level assignments were not consistently possible for all taxa because not all brachyurans species have been studied in detail. As in Klompmaker et al. (2014), both branchial sides needed to be preserved to confirm that specimens were not infested, whereas this was not a requirement for specimens with *Kanthyloma*. Our results show that there is no significant relationship between taxon abundance and infestation percentage on both the genus- and family-levels (Fig. 5). Similar results apply when genus-level data is split into Anomura (n = 6; r = −0.28, two-tailed t-test p = 0.59) and Brachyura (n = 10; r = −0.09, two-tailed t-test p = 0.80). Sample size is not adequate for such analyses on the family-level. Decapods from the Ernstbrunn Limestone assemblage were, however, not all collected at the same location or stratigraphic level (Bachmayer 1945). The precise locality and stratigraphic position are not known for all Ernstbrunn specimens. In situ collected assemblages from a single bed and small spatial scale with a relatively high proportion of parasitized specimens would be needed to further test the relationship between abundance and prevalence. Brachyurans and galatheoids from the Tithonian Štramberk Limestone in the active Kotouč Quarry (Czech Republic, Fraaije et al. 2013) may be mostsuitable because decapods from this limestone are occasionally infested (Houša 1963; pers. obs. AAK, CMR). Due to the generally low prevalence of epicarideans within large, diverse decapod assemblages today (Williams and Boyko 2012, and references therein), modern assemblages may not be suitable for this purpose.

**Figure 5.**
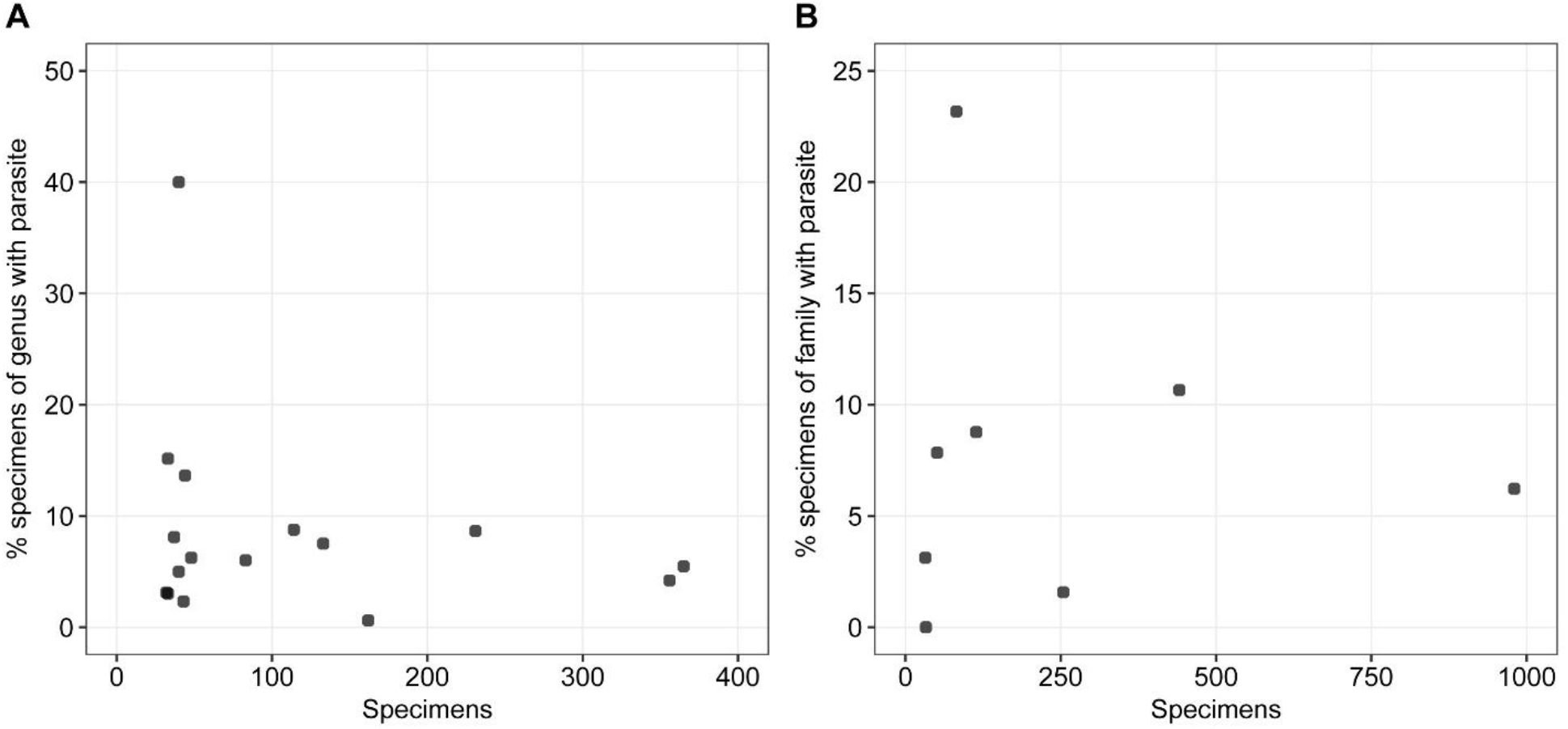
The percentage of parasitized specimens per taxon versus taxon abundance for decapods from the Ernstbrunn Limestone assemblage (Late Jurassic, Tithonian, Austria). A. Genera, n = 16; r = −0.23, two-tailed t-test p = 0.39. B. Families, n = 8; r = −0.04, two-tailed t-test p = 0.93. Minimum number of specimens per taxon = 30.

### 2.4. Host preference

Host preference is likely to result in non-random patterns of parasite prevalence. Some generalizations have been made concerning epicaridean parasites: Markham (1986) mentioned that host-species specificity appears to be rare among Bopyriodea; McDermott (1991) remarked that most branchial bopyroids infest particular crab families; and Boyko and Williams (2009) noted that some bopyroids infested multiple paguroid hosts, whereas others have been found only on one species thus far. Data in systematic studies on epicarideans echo variability in host specificity: An et al. (2009: table 1) showed that most *Progebiophilus* spp. were restricted to one thalassinidean host species and both An et al. (2015) and Boyko et al. (2017) noted epicaridean species infecting only a single host species, whereas other epicaridean species were found on multiple, often related host species. For carid shrimp from Florida, each bopyrid species was found only on one host genus (Briggs et al. 2017). The fact that multiple species from the same genus or family can be infested by the same parasite species does not imply a lack of preference because parasitized hosts can have very different infestation percentages (e.g., Owens and Glazebrook 1985; González and Acuña 2004; Brockerhoff 2004).

Host specificity of individual parasite species is impossible to address because epicarideans tend not to preserve as fossils due to their low preservation potential (Klompmaker et al. 2017; Fig. 2), but it is possible to assess host preferences for all epicaridean-induced swellings combined. Very little is known thus far. From the Late Cretaceous (Campanian-Maastrichtian) of Greenland, species-level infestation percentages have been reported from a lobster-brachyuran assemblage preserved in siliciclastic rocks, primarily in shales (Collins and Wienberg Rasmussen 1992; Wienberg Rasmussen et al. 2008). The raninoid crab species *Macroacaena rosenkrantzi* (Collins and Wienberg Rasmussen, 1992) was infested more frequently than the congeneric *M. succedana* (Collins and Wienberg Rasmussen, 1992) (73/1295 = 5.6% vs a few out of 193 =< 2%). Species-level data is also known from the mid-Cretaceous reefal limestones (late Albian) from Spain (Klompmaker et al. 2014). The galatheoid *Eomunidopsis navarrensis* (Van Straelen, 1940) was heavily infested (21/174 = 12.1%) relative to other common species (< 5%).

To evaluate host preference for the Jurassic for the first time, the Tithonian Ernstbrunn Limestone decapod assemblage was used again and the same specimen selection criteria as above were used with specimens determined to the genus-level. We studied host preference in a more quantitative framework than was done previously, testing host preference against a null model as described in Smith et al. (2018). To test for host preference in the entire assemblage, the observed number of infested specimens per taxon was compared to the expected number in the null model (i.e., random distribution) using the Bray-Curtis index of dissimilarity based on 10,000 iterations. Subsequently, preference for individual host genera are assessed using Manly’s alpha to identify which genera are more and less frequently infested than would be expected by chance. The results indicate that evidence for *Kanthyloma crusta* is distributed non-randomly across the genera because only 0.05% of the simulated Bray-Curtis dissimilarity values is greater than the observed Bray-Curtis dissimilarity value (Fig. 6A). The only taxon that is preferentially infested is the moderately common genus *Discutiolira* Robins et al., 2016, whereas *Longodromites* Patrulius, 1959, *Prosopon* von Meyer, 1835, *Lecythocaris* von Meyer, 1858, *Gastrodorus* von Meyer, 1864, *Abyssophthalmus* Schweitzer and Feldmann, 2009b, *Glaessneropsis* Patrulius, 1959, and *Nodoprosopon* Beurlen, 1928, are less frequently infested than expected by chance (Fig. 6B). To compare the results to the mid-Cretaceous Koskobilo assemblage (Klompmaker et al. 2014: table 1), the same tests were run on the genus- and species-levels from that assemblage. Bray-Curtis dissimilarity analyses indicate that *Kanthyloma crusta* is distributed non-randomly (insets Fig. 6C, D). The very abundant *Eomunidopsis* Vía Boada, 1981, and the less common *Faksecarcinus* Schweitzer et al., 2012, are preferential host taxa, while *Eodromites* Patrulius, 1959, appears avoided by epicarideans (Fig. 6C). On the species-level, *Eomunidopsis navarrensis* and *E. aldoirarensis* Klompmaker et al., 2012, are targeted, but *E. orobensis* (Ruiz de Gaona 1943), *Eodromites grandis* (von Meyer, 1857), *Distefania incerta* (Bell, 1863), and *Rathbunopon obesum* (Van Straelen, 1944) are less frequently infested than would be expected by chance (Fig. 6D).

**Figure 6.**
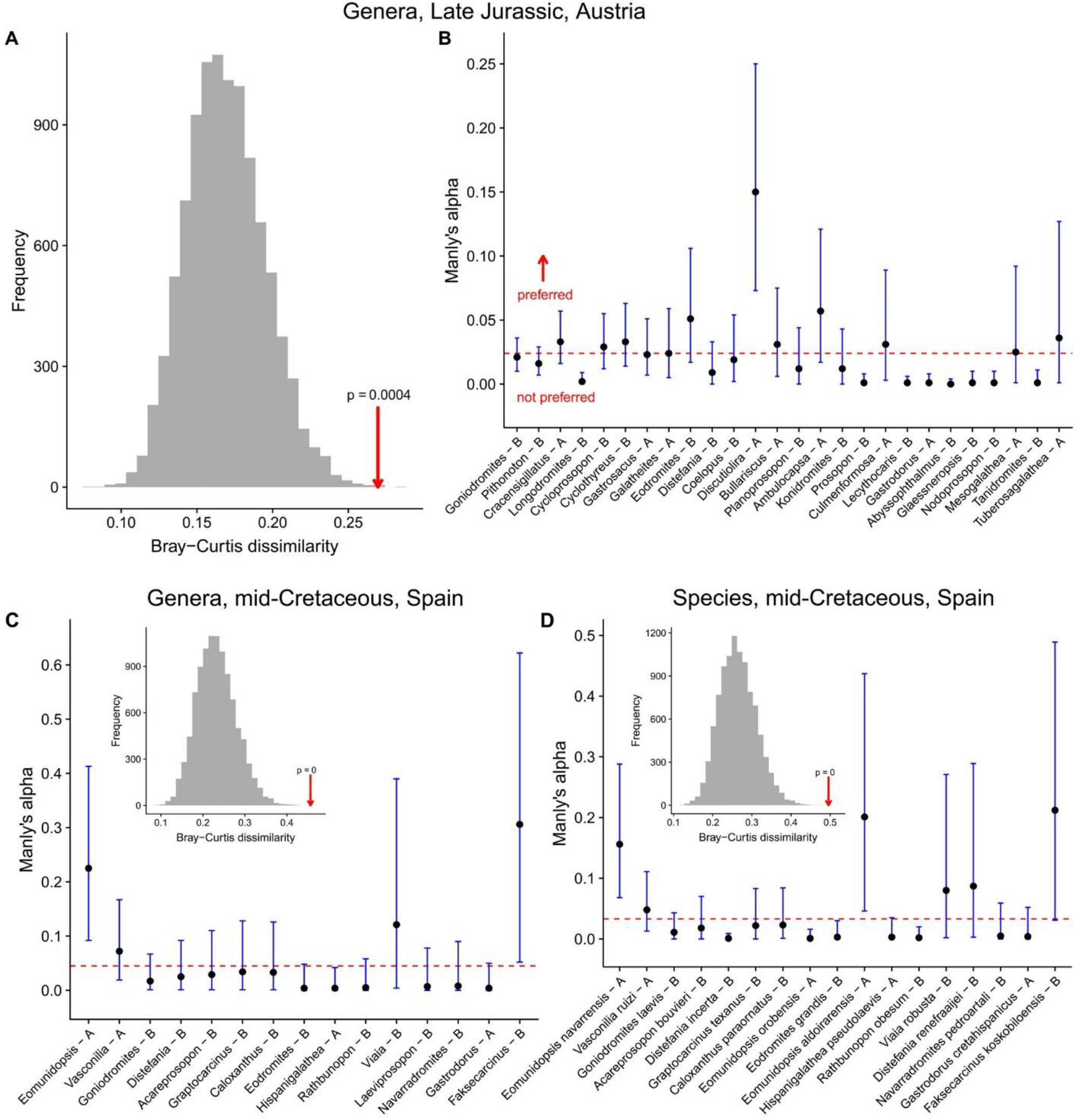
Parasite host preference analyses for decapods from the Ernstbrunn (Late Jurassic, Tithonian, Austria) and Koskobilo (mid-Cretaceous, late Albian, Spain) assemblages. A–B. Genera from Ernstbrunn, n = 42. C. Genera Koskobilo, n = 22. D. Species Koskobilo, n = 30. Histogram panels: grey histograms show randomized Bray-Curtis dissimilarity values assuming no host preference; red arrows indicate actual value for the assemblage. P-value is the chance of getting a higher value than the actual value. Panels with dots and error bars: Manly’s alpha values with 95% confidence intervals for brachyuran (B) and anomuran (A) taxa, ordered by decreasing abundance. Dashed line is the expected alpha value for each taxon (1 / n), assuming no host preference. Only taxa with at least ten specimens are shown.

Host specificities can be compared across assemblages by different methods. Basic host specificity considers the number of taxa infested (Poulin et al. 2011), which is higher for Ernstbrunn (22 genera) than for Koskobilo (9 genera). However, the Ernstbrunn Limestone assemblage contains more genera (42 vs 22). To remedy this issue, the percentage of genera that is parasitized can be compared. This method yields a higher percentage for Ernstbrunn (52%) than for vs Koskobilo (41%). A caveat here is that many of the non-infested genera are represented by a low number of specimens. Other indices compare the prevalence per host taxon or structural specificity (Poulin et al. 2011). Rohde’s modified index takes into account the percentage of parasitized specimens per taxon and corrects for a different number of host genera (Rohde 1980; Rohde and Rohde 2005). This index ranges from 0 to 1 with a higher value implying a higher level of host specificity among the infested hosts. Using genus-level data, the mid-Cretaceous Koskobilo assemblage yield a higher host specificity value than the Late Jurassic Ernstbrunn Limestone assemblage (0.49 vs 0.39). Another specificity measure combines prevalence and phylogenetic distance to compare host specificity (Poulin and Mouillot 2005), but the phylogenetic structures of the Ernstbrunn and Koskobilo assemblages differ substantially, rendering any comparisons using this method uninformative.

### 2.5. Size of parasitized versus non-parasitized specimens

Parasitism by epicaridean isopods poses a metabolic drain on the decapod host because female parasite individuals feed on host hemolymph and ovarian fluids after piercing the host’s cuticle (Bursey 1978; Lester 2005). For example, a bopyrid parasite can consume up to 25% (Walker 1977) or up to 10% (Anderson 1977) of hemolymph volume per day, for a species of carid shrimp. Consequently, a lower growth rate leading to smaller average and maximum sizes within parasitized specimens may be expected. Multiple studies have indeed shown that modern parasitized decapods are smaller on average than uninfected specimens of the same taxon (e.g., Roccatagliata and Lovrich 1999, for a lithodid anomuran; González and Acuña 2004, for a galatheoid; Petrić et al. 2010, for a galatheoid), but other studies found that infested individuals are of similar size (e.g., Mantelatto and Miranda 2010, for porcellanid anomuran) or even larger (e.g., McDermott 1991, for a brachyuran; Lee et al. 2016, for a carid shrimp). For maximum size, many examples exist in which the non-parasitized individuals were larger (O’Brien and Van Wyk 1985, and references; Roccatagliata and Lovrich 1999), but one study showed no difference (O’Brien and Van Wyk 1985).

No study has compared body sizes of parasitized versus non-parasitized fossil decapods of the same taxon thus far. To that end, we examined two species of the Ernstbrunn Limestone assemblage introduced above. Both the galatheoid *Cracensigillatus acutirostris* (Moericke, 1889) and the brachyuran *Goniodromites bidentatus* Reuss, 1858, are abundant species in this assemblage, specimens of these taxa are fairly well-preserved, and multiple specimens are infested, making them suitable target species. Using digital calipers, the maximum width (without spines and without the additional width due to the parasitic swelling) and maximum length (without the long, often incomplete rostrum for *C. acutirostris*) was measured for all infested specimens where possible. For non-parasitized specimens, the same measurements were taken for 43 and 36 randomly chosen individuals of *C. acutirostris* and *G. bidentatus*, respectively. Length (L), width (W), and the geometric mean of length and width (√(L×W)) are used to compare the sizes of parasitized and non-parasitized specimens for both species using a Mann-Whitney test and a significance level of 5%.

For the geometric mean (Fig. 7), the median size is significantly larger for parasitized specimens of *C. acutirostris* (p = 0.008), while the median appears also larger for *G. bidentatus*, but not significantly so (p = 0.057). Results are similar for the length (p = 0.030 and p = 0.060, resp.), but the median widths are statistically larger for parasitized specimens of both species (p = 0.002 and p = 0.018, resp.). Larger size classes contain a higher proportion of infested specimens, although sample size is limited for some size classes (Fig. 7).

**Figure 7.**
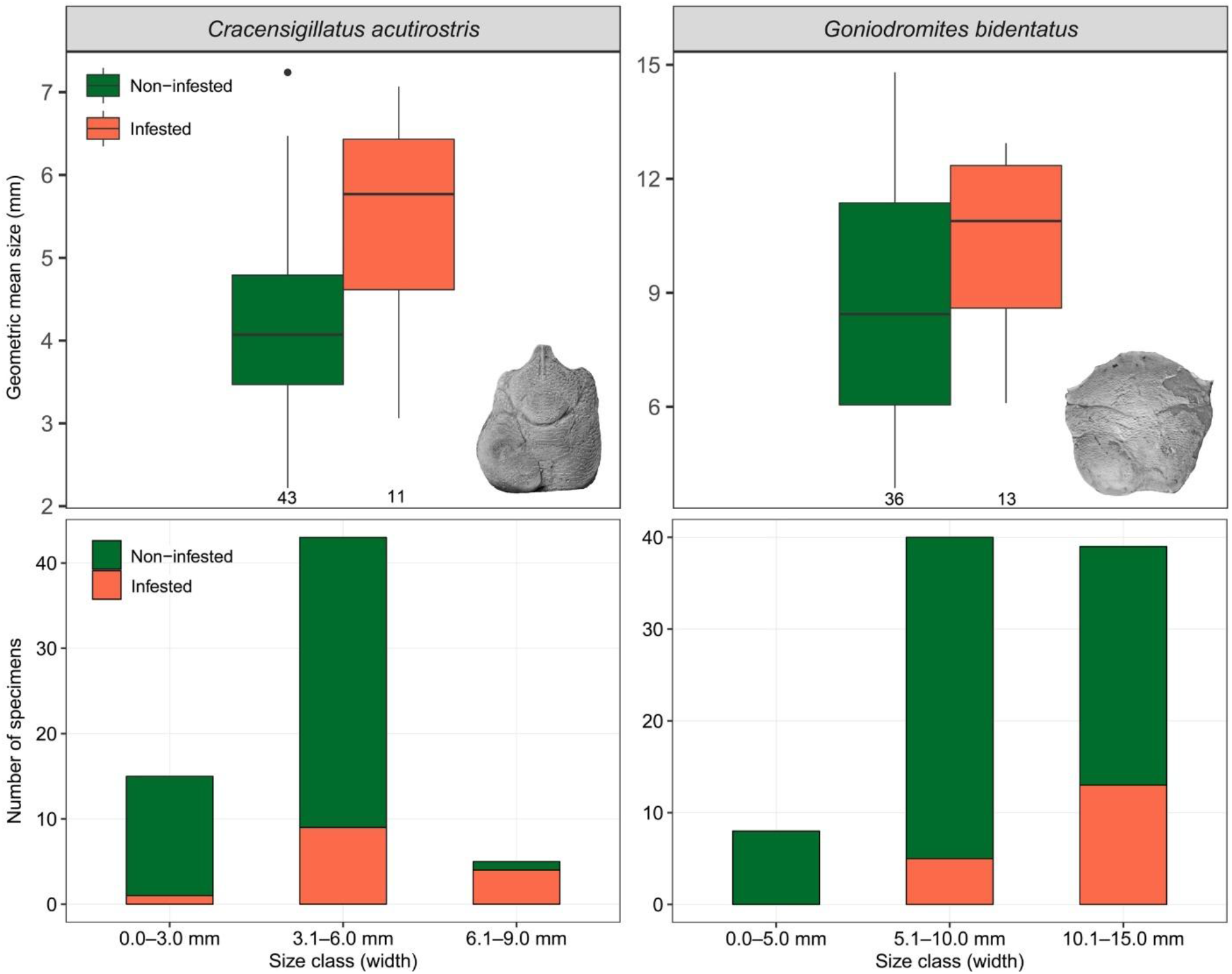
Upper panel: boxplots of geometric mean sizes of non-infested vs infested specimens of two decapod species from the Ernstbrunn Limestone assemblage (Late Jurassic, Tithonian, Austria). Geometric mean size = √ [maximum carapace length (excluding rostrum for *C. acutirostris*) × maximum carapace width excluding spines]. Infested galatheoid *Cracensigillatus acutirostris* (photo = NHMW 2007z0149/0315, modified from Robins et al. 2013: fig. 16.12): Mann-Whitney p = 0.008. Infested brachyuran *Goniodromites bidentatus* (photo = NHMW 2017/0089/0012): Mann-Whitney p = 0.057. Lower panel: per size class stacked bar diagrams of non-infested and infested specimens for the two decapod species. Maximum width is used to maximize sample size.

For maximum size, the samples of non-infested specimens contain the largest specimen for both species (see Fig. 7), but non-infested specimens are more numerous than infested specimens in both cases. To test the effect of unequal sample size on the maximum size, bootstrap analyses without replacement were carried out. The maximum size was determined 10,000 times from random sampling without replacement from the pool of non-infested specimens with the total number of infested specimens for that species as the number of specimens to be sampled from the pool of non-infested specimens. The distribution of maximum sizes was then compared to the maximum size of the infested sample for both species, with a p-value representing the chance of getting a larger non-parasitized specimen than the largest parasitized specimen here. For both species, the maximum sizes of the infested specimens fall well within the distribution of bootstrapped maximum sizes for non-infested specimens (p > 0.05, Fig. 8), indicating that the maximum sizes of parasitized and non-parasitized taxa do not differ statistically.

**Figure 8.**
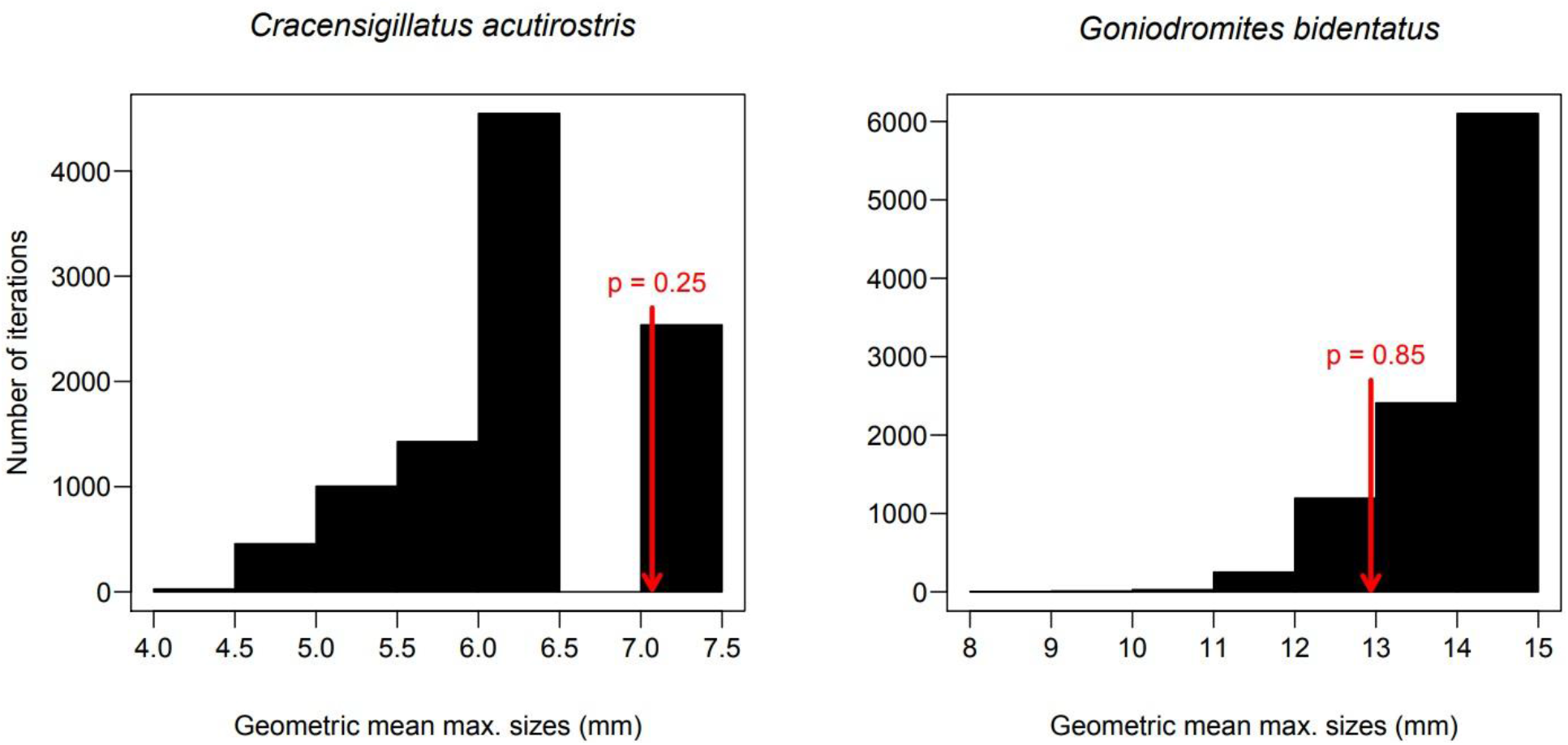
Comparison of the maximum size of a parasitized specimen versus bootstrapped maximum sizes of non-parasitized specimens for two decapod species from the Ernstbrunn Limestone assemblage (Late Jurassic, Tithonian, Austria). Bootstrapping without replacement (number of iterations = 10,000) was performed because the sample size of non-parasitized specimens was larger than that of parasitized specimens in both cases (sample sizes as in previous figure). Red arrow represents the maximum size of parasitized specimens within the sample. P-value is the chance of getting a larger non-parasitized specimen than the largest parasitized specimen.

How can the larger median size of infested specimens be explained for both species? We will discuss the following hypotheses: (1) swellings in small specimens were not recognized, (2) parasitized specimens represent a different assemblage consisting of larger specimens on average, (3) small specimens with a swelling have a relatively low preservation potential compared to similar-sized specimens that are not infested and/or small specimens have a lower preservation potential than large infested carapaces, (4) the larger swollen specimens represented specimens of which the parasite was lost at an earlier stage but the swelling remained present, (5) parasites have an equal probability of infesting their host at any point during the life of the host, (6) parasites are more likely to infest larger hosts, (7) complete or partial castration of the host leads to increased growth and/or longevity of parasitized hosts, (8) juvenile hosts have a higher probability of dying than adult infested specimens, (9) parasites selected for the larger sex within species, and (10) a combination of several factors.

It is unlikely that swellings in small specimens were not noted or were too small to be distinguished from possible post-mortem deformations. We (AAK, CMR) studied specimens for swellings with 10x magnification hand lenses and from different angles in case of doubt.

Not all specimens from the Ernstbrunn Limestone assemblage were collected in the same locality and possibly the same stratigraphic level (see above), which may affect host size and parasite prevalence (e.g., McDermott 1991). Bachmayer (1948) reported on parasitic swellings in nine individuals across three species, six from Dörfles I and three from Dörfles Werk II, and Bachmayer (1964) showed another specimen from Dörfles Werk II with *Kanthyloma*. Analyzing the specimens per locality is not possible because the exact locality is unknown for most specimens. Given that Dörfles I and Dörfles Werk II are only ~100 m apart (Bachmayer 1945; Schneider et al. 2013), the hypothesis that the parasitized specimens represent a very different assemblage consisting of larger individuals appears unlikely.

No experiments have explored whether small corpses and/or molts containing a swelling have a lower preservation potential than equally-sized specimens without a swelling. However, a galatheoid host cuticle was much thicker on the swelling due to a thicker epidermis/connective tissue layer relative to a non-infested conspecific, but the calcified layers were of equal thickness (Bursey 1978). For non-infested specimens, the decay rate of small specimens is not markedly different than in large specimens for most marine arthropods, including brachyurans (Klompmaker et al. 2017). Thus, a relatively low preservation potential for small specimens exhibiting *Kanthyloma* is unlikely to explain the larger size of infested specimens.

Not all decapods exhibiting a swollen branchial region harbor an isopod parasite permanently, with the isopod leaving the branchial region prior to the death of the host. Such occurrences, however, cannot be identified in fossil specimens because the preservation potential for epicaridean isopods is low (Klompmaker et al. 2017; Fig. 2). Decapods can lose the parasite during the molting phase either after the death of the parasite or when the living parasite gets dislodged (e.g., Van Wyk 1982; Anderson 1990; Somers and Kirkwood 1991; Cash and Bauer 1993; Roccatagliata and Lovrich 1999), but the swelling in the carapace can remain present even after a new molting phase for at least one molting cycle for a porcellanid or even four to five cycles for a carid shrimp (Van Wyk 1982). Such empty swellings can dominate large size classes of infested specimens (Van Wyk 1982; Roccatagliata and Lovrich 1999), but they make up only a small portion of all specimens in those size classes (Van Wyk 1982; Cash and Bauer 1993; Roccatagliata and Lovrich 1999). Conversely, no marked increase of empty swellings with carapace size were found for a galatheoid (Wenner and Windsor 1979) and a porcellanid (Oliveira and Masunari 1998). Thus, carapaces with these “ghost parasites” are unlikely to result in a substantial increase in the median sizes of fossil specimens with swellings.

Regarding the timing of infestation, infestation of many modern decapods by bopyroids is suggested to occur almost exclusively in juvenile hosts based on (1) a positive and often significant relationship between parasite size and host size (e.g., Allen 1966; Beck 1980; Abu-Hakima 1984; Cash and Bauer 1993; Oliveira and Masunari 1998; Roccatagliata and Lovrich 1999; González and Acuña 2004; Mantelatto and Miranda 2010; Román-Contreras and Romero-Rodríguez 2013; Baeza et al. 2018), (2) the observation that immature parasites are found in small hosts only (Van Wyk 1982; Oliveira and Masunari 1998, both for a porcellanids), and (3) small specimens are shown to be more readily infested than large specimens (Anderson 1990, for a carid shrimp). Consequently, swellings should be visible early on in the life of infested decapod hosts and, all else being equal, the proportion of specimens that is infested would remain about the same or decrease due to the adverse effects of parasitism on the host (see above) as host size increases. Such patterns are indeed shown in papers that also report on parasite prevalence per size class (e.g., Oliveira and Masunari 1998; Roccatagliata and Lovrich 1999; González and Acuña 2004; Mantelatto and Miranda 2010; but see Román-Contreras and Romero-Rodríguez 2013). On the other hand, epicarideans can also infest their host throughout host ontogeny as shown by a relatively weak or lack of correlation between parasite size (including immature specimens) and host size (Brockerhoff 2004; Smith et al. 2008; Griffen 2009; Rasch and Bauer 2015). Research using only mature female parasites to test for an expected correlation between parasite size and host size (e.g., McDermott 1991; Roccatagliata and Lovrich 1999; Jordá and Roccatagliata 2002) should not be taken into account in this case because all epicarideans should be used to assess the timing of infestation. All else being equal, an increase in prevalence with host size is expected when infestation takes place throughout host ontogeny or primarily in larger size classes. As this pattern matches the size-prevalence results herein (Fig. 7), infestation throughout ontogeny or preferential infection of larger specimens rather than solely in juveniles can explain the larger median size of infested individuals. This hypothesis is consistent with the lack of epicarideans in small host size classes in some studies (Mantelatto and Miranda 2010; Lee et al. 2016).

Castration may provide another explanation for the larger median size of infested specimens, if the energy that would have been spent on host reproduction is directed toward growth of the host specimen (cf. Poulin 2011: fig. 5.7) instead of to the growth of the parasite and the parasite’s reproductive efforts. Host gigantism may be (1) a strategy of the parasite because parasite fecundity is positively linked to host size and host survival may perhaps be improved or (2) an adaptation of the host to counterbalance the negative impact of parasitism (Poulin 2011). Parasite fecundity is often positively correlated with some measure of bopyroid parasite size (e.g., Jay 1989; McDermott 2002; Romero-Rodríguez and Román-Contreras 2008; Román-Contreras and Romero-Rodríguez 2013; Cericola and Williams 2015; Baeza et al. 2018) or host size (Beck 1980); thus, a larger host would be beneficial to the parasite population. However, not all epicarideans castrate their host and host growth rates are inhibited relative to non-infested specimens for many decapods when the epicaridean becomes mature, perhaps because more energy is diverted to the parasite at that point (O’Brien and Van Wyk 1985, and references; Romero-Rodríguez et al. 2016). Castrated hosts that outlive non-parasitized conspecifics would also result in relatively large parasitized specimens. However, at least some species live shorter lives (Romero-Rodríguez et al. 2016, for a carid shrimp) and the multiple negative effects of parasitism (see above) render the castration hypothesis unlikely.

A larger median size of infested individuals may be explained by young hosts having a higher probability of dying than older infested specimens, particularly when a branchial swelling has yet to form in small specimens. For those that do survive, a swelling may appear only in larger size classes. High mortality of primarily young hosts infested by a bopyrid is known for a carid shrimp during the one-to-two-week endoparasitic stage before becoming ectoparasitic in the gill chamber (Anderson 1990). Mortality of infested vs non-infested individuals was equal after five weeks (Anderson 1990), but specimen sizes were not provided. Others also hinted at a high mortality of young, infested individuals (Roccatagliata and Lovrich 1999; Lee et al. 2016). As little is known about how much time it takes for an epicaridean to cause a swelling in small and large hosts, it is difficult to further evaluate this hypothesis.

Selection of the larger sex of a species, which would have a positive effect on parasite fecundity (see above), could result in a larger median size for infested specimens. For a galatheoid, Petrić et al. (2010) found that the larger male specimens were infected more frequently despite being less common. Our analysis of Petrić et al’s data indicates that males are significantly more infested than females (χ^2^ = 4.7, p = 0.03). Beck (1979) mentioned several more examples of bopyroids infecting the larger sex, although results were not always consistent and sexes do not always differ in size (e.g., McDermott 1991). If both fossil *Cracensigillatus acutirostris* and *Goniodromites bidentatus* had differently-sized sexes, it is possible that the larger sex was selected by epicarideans. Unfortunately, sex determination is not possible for the fossil specimens because only dorsal carapaces were available.

In sum, despite a possible slower growth rate for infested specimens, infestation throughout ontogeny rather than exclusively in young individuals and, possibly, selection for the larger sex appear the most likely explanations for the larger median sizes of infested specimens of *C. acutirostris* and *G. bidentatus*. This study adds to the rare examples from the fossil record testing the relative sizes of infested and non-infested specimens. Other examples include Paleozoic crinoids parasitized by platyceratid gastropods that were either larger (Baumiller and Gahn 2018) or smaller (Rollins and Brezinski 1988; Gahn and Baumiller 2003) than non-parasitized crinoids, and Holocene and possibly Pleistocene bivalves infested by trematodes that were larger than non-infested specimens (Ruiz and Lindberg 1989; Huntley and Scarponi 2012).

## 3. Rhizocephalan barnacles in decapod crustaceans

Rhizocephalan barnacles infest modern decapods and castrate their host, cause feminization of male individuals, and reduce host growth rates (e.g., Reinhard 1956; O’Brien and Van Wyk 1985; Takahashi and Matsuura 1994; Høeg et al. 2005; Nagler et al. 2017b). Feminization of male individuals is seen in true crabs (e.g., Reinhard 1956), but not in squat lobsters (Boyko and Williams 2011). The fossil record of evidence for rhizocephalans parasitizing decapods has been discussed in detail recently (Klompmaker and Boxshall 2015). In short, feminization of male crab specimens from the Late Cretaceous of South Dakota, USA (Bishop 1974; Bishop 1983b; Jones 2013) and the Miocene of New Zealand (Feldmann 1998) is known, with the latter specifically attributed to Rhizocephala. However, epicarideans may also cause feminization of male specimens (e.g., Reinhard 1956; Rasmussen 1973; O’Brien and Van Wyk 1985). No new records of feminized fossil decapods have been reported since 2015, but a 5 mm large nauplius larva, perhaps close to Rhizocephala, was found in the Late Jurassic of Germany (Nagler et al. 2017a). Collections of decapods with ventral sides and appendages preserved would be suitable for additional studies on this type parasitism. Anomura have been suggested as the ancestral hosts of Rhizocephala because four basal rhizocephalans all infested anomurans and an ancestral host reconstruction also pointed toward anomurans (Glenner and Hebsgaard 2006; Scholtz et al. 2009).

## 4. Ciliates on ostracods

Phosphatized stalked peritrichid ciliates (Ciliophora) have been found attached to ostracods within an Early Triassic ammonoid from Svalbard (Spitsbergen) (Weitschat and Guhl 1994). The specimens were up to 0.2 mm long and attached to the inner part of the shell and on the epipodal appendages. These specimens were not considered parasitism in the strict sense because the specimens were filter feeding rather than feeding directly on the host (Klompmaker and Boxshall 2015). More specimens may only be found in Konservat-Lagerstätten.

## 5. “Pentastomids” on ostracods

Four 1–4 mm long specimens identified as Pentastomida associated with a Silurian ostracod were reported from the Herefordshire Lagerstätte in England (Siveter et al. 2015). Whether these specimens represent true pentastomids was called into question (De Baets et al. 2015; De Baets and Littlewood 2015; Klompmaker and Boxshall 2015). A re-evaluation rejects a pentastomid affinity because the snout and trunk are in different planes, unlike for true pentastomids in which they are in the same plane; the paired limbs are proportionally longer than in extant pentastomids; and no apical hooks were found, a feature characteristic of true pentastomids (Boxshall and Hayes in press). Regardless of the taxonomic identity, two specimens occurred jointly on the carapace, suggesting that these specimens may not have been nutritionally dependent on the ostracod (not parasitism in the strict sense), but two others were found within the ostracod at the position of the gills near eggs, and may have fed on eggs (parasitism). Specimens were identified as adults, suggesting that the ostracods may have served as the final host (Siveter et al. 2015).

## 6. Modern evidence with preservation potential

Many modern crustaceans serve as hosts of parasites sometime during the life cycle of parasites, but most are unlikely to be found in the fossil record (e.g., Klompmaker and Boxshall 2015). The reasons are: (1) the parasite does not leave a (recognizable) trace (e.g., Vannier and Abe 1993, for ostracod hosts); (2) the parasite is unlikely to be found as body fossils associated with the crustacean host because the parasite has a low preservation potential and/or is extremely small (e.g., Boxshall and Lincoln 1983, for tantulocarid parasites on other crustaceans); and/or (3) the body fossils of the host are unlikely to fossilize (e.g., Poinar et al. 2010, for amphipod hosts).

Some associations only found in modern ecosystems thus far have the potential to be found in the fossil record. First, the endoparasitic entoniscid isopods can cause swellings/asymmetries in the carapaces of decapod crustaceans including anomuran, brachyuran, and shrimp hosts (Miyashita 1941: p. 251; Shiino 1942: p. 62, 68; Shields and Kuris 1985: Fig. 1; Mushtaq et al. 2016: p. 1608). These swellings may be less prominent than those caused by the primarily ectoparasitic Bopyridae and Ionidae and not always at the same spot in the branchial chamber because entoniscids are found in the visceral cavity of their host and may even cause a swelling of the cardiac region (Adkison 1990; Williams and Boyko 2012: Fig. 1F; Mushtaq et al. 2016: Fig. 1). Although not all entoniscid cause malformations (McDermott 2009, for pinnotherid crabs), gentle bopyrid/ionid-induced swellings may be confused with swellings caused by entoniscid in fossil material. Thus, it is possible that some examples of *Kanthyloma crusta* ascribed to bopyrids/ionids are caused by entoniscids.

Second, Hosie (2008: fig. 12A) showed a barnacle *Smilium zancleanum* (Seguenza, 1876), with swelling up to 1 cm made by a cryptoniscoid isopod. The swellings are located in the muscular peduncle or at the base of the capitulum, where they can cause a disruption in the alignment of plates. Exceptional preservational circumstances are necessary for such swellings to preserve in the fossil record.

In addition to small (0.02–0.40 mm) hooks of likely platyhelminth (?Monogenea) origin found in fish, platyhelminth hooks have also been found in association with two crustacean specimens from the Late Devonian (Frasnian) of Latvia (Upeniece 2001; Upeniece 2011; De Baets et al. 2015). Two hooks were found near a specimen of Mysidacea (a member of Peracarida), whereas another hook was found near a clam shrimp (Conchostraca). Recent work has suggested that parasitism by platyhelminthes of these crustacean specimens is unlikely because of the large size of the hooks relative to the crustacean specimens (De Baets et al. 2015) and the rarity of hooks among the thousands of mysidacean specimens (Klompmaker and Boxshall 2015). Monogeneans parasitize nearly exclusively fish today, but another group of platyhelminthes, Cestoda, also uses hooks for attachment to crustacean host, including their larvae (e.g., Xylander 2005). Cestodes have been recorded from modern decapods such as true crabs (Dollfus 1976; Torchin et al. 2001), hermit crabs (McDermott et al. 2010), and shrimps (Dollfus 1976; Georgiev et al. 2007). Cestode hooks are mineralized (e.g., Collin 1968; Ambrosio et al. 2003). Thus, minute platyhelminth hooks may be found associated with fossil crustaceans in the right preservational settings in the future.

## Acknowledgements

AAK and CMR thank Thomas Nichterl and Andrea Krapf (both Naturhistorisches Museum Wien, Austria) for help with handling loans, and Andreas Kroh (Naturhistorisches Museum Wien) for facilitating a wonderful research stay in Vienna in 2017. René Fraaije (Oertijdmuseum, The Netherlands) made his fossil decapod collection available for study. This research was supported by a Paleontological Society Arthur J. Boucot research grant to AAK and, in part, by a Palaeontological Association research grant (PA-RG201401) to AAK.

## References

Abu-Hakima R (1984) Preliminary observations on the effects of *Epipenaeon elegans* Chopra (Isopoda; Bopyridae) on reproduction of *Penaeus semisulcatus* de Haan (Decapoda; Penaeidae). International Journal of Invertebrate Reproduction and Development 7:51–62. doi: 10.1080/01688170.1984.10510071

Adkison DL (1990) A review of the Entoniscinae (Isopoda: Epicaridea: Entoniscidae). Dissertation, Tulane University

Allen JA (1966) Notes on the relationship of the bopyrid parasite *Hemiarthrus abdominalis* (Krøyer) with its hosts. Crustaceana 10:1–6. doi: 10.1163/156854066X00018

Ambrosio J, Reynoso-Ducoing O, Hernández-Sanchez H, Correa-Piña D, González-Malerva L, Cruz-Rivera M, Flisser A (2003) Actin expression in *Taenia solium* cysticerci (cestoda): tisular distribution and detection of isoforms. Cell Biology International 27:727–733. doi: 10.1016/S1065-6995(03)00142-2

An J, Boyko CB, Li X (2015) A review of bopyrids (Crustacea: Isopoda: Bopyridae) parasitic on caridean shrimps (Crustacea: Decapoda: Caridea) from China. Bulletin of the American Museum of Natural History 399:1–85. doi: 10.1206/amnb-921-00-01.1

An J, Williams JD, Yu H (2009) The Bopyridae (Crustacea: Isopoda) parasitic on thalassinideans (Crustacea: Decapoda) from China. Proceedings of the Biological Society of Washington 122:225–246. doi: 10.2988/08-26.1

Anderson G (1977) The effects of parasitism on energy flow through laboratory shrimp populations. Marine Biology 42:239–251. doi: 10.1007/BF00397748

Anderson G (1990) Postinfection mortality of *Palaemonetes* ssp. (Decapoda: Palaemonidae) following experimental exposure to the bopyrid isopod *Probopyrus pandalicola* (Packard)(Isopoda: Epicaridea). Journal of Crustacean Biology 10:284–292

Anderson G (1975) Metabolic response of the caridean shrimp *Palaemonetes pugio* to infection by the adult epibranchial isopod parasite *Probopyrus pandalicola*. Comparative Biochemistry and Physiology Part A: Physiology 52:201–207. doi: 10.1016/S0300-9629(75)80153-8

Arneberg P, Skorping A, Grenfell B, Read AF (1998) Host densities as determinants of abundance in parasite communities. Proceedings of the Royal Society B: Biological Sciences 265:1283–1289. doi: 10.1098/rspb.1998.0431

Bachmayer F (1948) Pathogene Wucherungen bei jurassischen Dekapoden. Sitzungsberichte der Österreichischen Akademie der Wissenschaften, Mathematisch-naturwissenschaftliche Klasse, Abteilung I 157(6–10):263–266

Bachmayer F (1945) Die Crustaceen aus dem Ernstbrunner Kalk der Jura-Klippenzone zwischen Donau und Thaya. Jahrbuch der Geologischen Bundesanstalt 1945:35–43

Baeza JA, Steedman S, Prakash S, Liu X, Bortolini JL, Dickson M, Behringer DC (2018) Mating system and reproductive performance in the isopod *Parabopyrella lata*, a parasitic castrator of the ‘peppermint’ shrimp *Lysmata boggessi*. Marine Biology 165:41. doi: 10.1007/s00227-018-3297-z

Bass CS, Weis JS (1999) Behavioral changes in the grass shrimp, *Palaemonetes pugio* (Holthuis), induced by the parasitic isopod, *Probopyrus pandalicola* (Packard). Journal of Experimental Marine Biology and Ecology 241:223–233

Baumiller TK, Gahn FJ (2018) The nature of the platyceratid–crinoid association as revealed by cross-sectional data from the Carboniferous of Alabama (USA). Swiss Journal of Palaeontology 137:177–187. doi: 10.1007/s13358-018-0167-8

Beck JT (1979) Population interactions between a parasitic castrator, *Probopyrus pandalicola* (Isopoda: Bopyridae), and one of its freshwater shrimp hosts, *Palaemonetes paludosus* (Decapoda: Caridea). Parasitology 79:431–449. doi: 10.1017/S003118200005383X

Beck JT (1980) Life history relationships between the bopyrid isopod *Probopyrus pandalicola* and of its freshwater shrimp hosts *Palaemonetes paludosus*. American Midland Naturalist 104:135–154. doi: 10.2307/2424966

Beurlen K (1928) Die Decapoden des schwäbischen Jura mit Ausnahme der aus den oberjurassischen Plattenkalken stammenden. Palaeontographica 70:115–278, pls. 6–8

Bishop GA (1974) A sexually aberrant crab (*Dakoticancer overanus* Rathbun, 1917) from the upper cretaceous Pierre shale of South Dakota. Crustaceana 26:212–218. doi: 10.1163/156854074X00578

Bishop GA (1983) A second sexually aberrant specimen of *Dakoticancer overanus* Rathbun, 1917, from the Upper Cretaceous *Dakoticancer* assemblage, Pierre Shale, South Dakota (Decapoda, Brachyura). Crustaceana 44:23–26. doi: 10.1163/156854083X00028

Bosc LAG (1802) Histoire naturelle des vers, contenant leur description et leurs moeurs, avec figures dessinees d’après nature, volumes 1–3. Deterville, Paris

Boxshall G, Lester R, Grygier M, Høeg J, Glenner H, Shields J, Lützen J (2005) Crustacean parasites. In: Rohde K (ed) Marine Parasitology. CSIRO Publishing, Collingwood, pp 123–169

Boxshall GA, Hayes P (In press) Biodiversity and taxonomy of the parasitic Crustacea. In: Smit N, Bruce N (eds) Parasitic Crustacea. Springer

Boxshall GA, Lincoln RJ (1983) Tantulocarida, a new class of Crustacea ectoparasitic on other crustaceans. Journal of Crustacean Biology 3:1–16. doi: 10.2307/1547849

Boyko CB, Moss J, Williams JD, Shields JD (2013) A molecular phylogeny of Bopyroidea and Cryptoniscoidea (Crustacea: Isopoda). Systematics and Biodiversity 11:495–506. doi: 10.1080/14772000.2013.865679

Boyko CB, Williams JD (2009) Crustacean parasites as phylogenetic indicators in decapod evolution. In: Martin JW, Crandall KA, Felder DL (eds) Decapod Crustacean Phylogenetics. CRC Press, Boca Raton, pp 197–220

Boyko CB, Williams JD (2011) Parasites and other symbionts of squat lobsters. In: Poore GCB, Ahyong ST, Taylor J (eds) The Biology of Squat Lobsters. CRC Press, Boca Raton, pp 271–295

Boyko CB, Williams JD, Shields JD (2017) Parasites (Isopoda: Epicaridea and Nematoda) from ghost and mud shrimp (Decapoda: Axiidea and Gebiidea) with descriptions of a new genus and a new species of bopyrid isopod and clarification of *Pseudione* Kossmann, 1881. Zootaxa 4365:251–301. doi: 10.11646/zootaxa.4365.3.1

Briggs SA, Blanar CA, Robblee MB, Boyko CB, Hirons AC (2017) Host abundance, sea-grass cover, and temperature predict infection rates of parasitic isopods (Bopyridae) on caridean shrimp. Journal of Parasitology 103:653–662. doi: 10.1645/16-126

Brinton BA, Curran MC (2015) The effects of the parasite *Probopyrus pandalicola* (Packard, 1879) (Isopoda, Bopyridae) on the behavior, transparent camouflage, and predators of *Palaemonetes pugio* Holthuis, 1949 (Decapoda, Palaemonidae). Crustaceana 88:1265–1281. doi: 10.1163/15685403-00003501

Brockerhoff AM (2004) Occurrence of the internal parasite *Portunion* sp. (Isopoda: Entoniscidae) and its effect on reproduction in intertidal crabs (Decapoda: Grapsidae) from New Zealand. Journal of Parasitology 90:1338–1344. doi: 10.1645/GE-295R

Bursey CR (1978) Histopathology of the parasitization of *Munida iris* (Decapoda: Galatheidae) by *Munidion irritans* (Isopoda: Bopyridae). Bulletin of Marine Science 28:566–570

Calado R, Vitorino A, Dinis MT (2006) Bopyrid isopods do not castrate the simultaneously hermaphroditic shrimp *Lysmata amboinensis* (Decapoda: Hippolytidae). Diseases of Aquatic Organisms 73:73–76. doi: 10.3354/dao073073

Cash CE, Bauer RT (1993) Adaptations of the branchial ectoparasite *Probopyrus pandalicola* (Isopoda: Bopyridae) for survival and reproduction related to ecdysis of the host, *Palaemonetes pugio* (Caridea: Palaemonidae). Journal of Crustacean Biology 13:111–124. doi: 10.1163/193724093X00480

Ceccon L, De Angeli A (2013) Segnalazione di decapodi eocenici infestati da parassiti isopodi (Epicaridea) (Vicenza, Italia settentrionale). Lavori Società veneziana di Scienze naturali 38:83–92

Cericola MJ, Williams JD (2015) Prevalence, reproduction and morphology of the parasitic isopod *Athelges takanoshimensis* Ishii, 1914 (Isopoda: Bopyridae) from Hong Kong hermit crabs. Marine Biology Research 11:236–252

Collin WK (1968) Electron microscope studies of the muscle and hook systems of hatched oncospheres of *Hymenolepis citelli* McLeod, 1933 (Cestoda: Cyclophyllidea). The Journal of Parasitology 54:74–88. doi: 10.2307/3276878

Collins JSH, Wienberg Rasmussen H (1992) Upper Cretaceous-Lower Tertiary decapod crustaceans from West Greenland. Bulletin fra Grønlands geologiske Undersøgelse 162:1–46

Combes C (2001) Parasitism. The ecology and evolution of intimate interactions. University of Chicago Press, Chicago

Corrêa LL, Oliveira Sousa EM, Flores Silva LV, Adriano EA, Brito Oliveira MS, Tavares-Dias M (2018) Histopathological alterations in gills of Amazonian shrimp *Macrobrachium amazonicum* parasitized by isopod *Probopyrus bithynis* (Bopyridae). Diseases of Aquatic Organisms 129:117–122. doi: 10.3354/dao03236

Cressey RF (1983) Crustaceans as parasites of other organisms. In: Bliss DE (ed) The biology of Crustacea, Vol. 6. Pathobiology. Academic Press, New York, pp 251–273

De Angeli A, Garassino A (2002) Galatheid, chirostylid and porcellanid decapods (Crustacea, Decapoda, Anomura) from the Eocene and Oligocene of Vicenza (N Italy). Memorie della Società Italiana di Scienze Naturali e del Museo Civico di Storia Naturale di Milano 30:1–40

De Baets K, Dentzien-Dias P, Upeniece I, Verneau O, Donoghue PCJ (2015) Constraining the deep origin of parasitic flatworms and host-interactions with fossil evidence. Advances in Parasitology 90:93–135. doi: 10.1016/bs.apar.2015.06.002

De Baets K, Littlewood DTJ (2015) The importance of fossils in understanding the evolution of parasites and their vectors. Advances in Parasitology 90:1–51. doi: 10.1016/bs.apar.2015.07.001

Dollfus R-P (1976) Énumération des Cestodes du plancton et des Invertébrés marins. 9e Contribution. Annales de Parasitologie 51:207–220

Feldmann RM (1998) Parasitic castration of the crab, *Tumidocarcinus giganteus* Glaessner, from the Miocene of New Zealand: Coevolution within the Crustacea. Journal of Paleontology 72:493–498. doi: 10.1017/S0022336000024264

Förster R (1969) Epökie, Entökie, Parasitismus und Regeneration bei fossilen Dekapoden. Mitteilungen aus der Bayerischen Staatssammlung für Paläontologie und historische Geologie 9:45–59

Fraaije RHB, Van Bakel BWM, Jagt JWM, Skupien P (2013) First record of paguroid anomurans (Crustacea) from the Tithonian-lower Berriasian of Štramberk, Moravia (Czech Republic). Neues Jahrbuch für Geologie und Paläontologie - Abhandlungen 269:251–259. doi: 10.1127/0077-7749/2013/0348

Gahn FJ, Baumiller TK (2003) Infestation of Middle Devonian (Givetian) camerate crinoids by platyceratid gastropods and its implications for the nature of their biotic interaction. Lethaia 36:71–82. doi: 10.1080/00241160310003072

Gemmellaro GG (1869) Studi paleontologici sulla fauna del calcare à Terebratula janitor del Nord di Sicilia. Part 1:11–18, pls. 2–3

Georgiev BB, Sánchez MI, Vasileva GP, Nikolov PN, Green AJ (2007) Cestode parasitism in invasive and native brine shrimps (*Artemia* spp.) as a possible factor promoting the rapid invasion of *A. franciscana* in the Mediterranean region. Parasitology Research 101:1647–1655. doi: 10.1007/s00436-007-0708-3

Glenner H, Hebsgaard MB (2006) Phylogeny and evolution of life history strategies of the parasitic barnacles (Crustacea, Cirripedia, Rhizocephala). Molecular Phylogenetics and Evolution 41:528–538. doi: 10.1016/j.ympev.2006.06.004

González MT, Acuña E (2004) Infestation by *Pseudione humboldtensis* (Bopyridae) in the squat lobsters *Cervimunida johni* and *Pleuroncodes monodon* (Galatheidae) off northern Chile. Journal of Crustacean Biology 24:618–624

Griffen BD (2009) Effects of a newly invasive parasite on the burrowing mud shrimp, a widespread ecosystem engineer. Marine Ecology Progress Series 391:73–83. doi: 10.3354/meps08228

Haug JT, Haug C, Nagler C (In press) Evolutionary history of crustaceans as parasites. In: De Baets K, Huntley JW (eds) The Evolution and Fossil Record of Parasitism. Springer, Dordrecht

Hernáez P, Martínez-Guerrero B, Anker A, Wehrtmann IS (2010) Fecundity and effects of bopyrid infestation on egg production in the Caribbean sponge-dwelling snapping shrimp *Synalpheus yano* (Decapoda: Alpheidae). Journal of the Marine Biological Association of the United Kingdom 90:691–698

Høeg JT, Glenner H, Shields JD (2005) Cirripedia Thoracica and Rhizocephala (barnacles). In: Rohde K (ed) Marine Parasitology. CSIRO Publishing, Wallingford, pp 154–165

Hosie AM (2008) Four new species and a new record of Cryptoniscoidea (Crustacea: Isopoda: Hemioniscidae and Crinoniscidae) parasitizing stalked barnacles from New Zealand. Zootaxa 1795:1–28

Houša V (1963) Parasites of Tithonian decapod crustaceans (Štramberk, Moravia). Sborník Ústředního ústavu geologického 28:101–114

Huntley JW, Scarponi D (2012) Evolutionary and ecological implications of trematode parasitism of modern and fossil northern Adriatic bivalves. Paleobiology 38:40–51. doi: 10.1666/10051.1

Hyžný M, Starzyk N, Robins CM, Kočová Veselská M (2015) Taxonomy and palaeoecology of a decapod crustacean assemblage from the Oxfordian of Stránská skála (Southern Moravia, Czech Republic). Bulletin of Geosciences 90:633–650. doi: 10.3140/bull.geosci.1559

Jay CV (1989) Prevalence, size and fecundity of the parasitic isopod *Argeia pugettensis* on its host shrimp *Crangon francisorum*. American Midland Naturalist 121:68–77

Jones A (2013) Population dynamics of Dakoticancer overanus from the Pierre Shale, South Dakota. MS Thesis, Kent State University

Jordá MT, Roccatagliata D (2002) Population dynamics of *Leidya distorta* (Isopoda: Bopyridae) infesting the fiddler crab *Uca uruguayensis* at the Río de la Plata Estuary, Argentina. Journal of Crustacean Biology 22:719–727. doi: 10.1163/20021975-99990286

Klompmaker AA, Feldmann RM, Robins CM, Schweitzer CE (2012) Peak diversity of Cretaceous galatheoids (Crustacea, Decapoda) from northern Spain. Cretaceous Research 36:125–145. doi: 10.1016/j.cretres.2012.03.003

Klompmaker AA, Artal P, Van Bakel BWM, Fraaije RHB, Jagt JWM (2014) Parasites in the fossil record: a Cretaceous fauna with isopod-infested decapod crustaceans, infestation patterns through time, and a new ichnotaxon. PLoS One 9:e92551. doi: 10.1371/journal.pone.0092551

Klompmaker AA, Boxshall GA (2015) Fossil crustaceans as parasites and hosts. Advances in Parasitology 90:233–289. doi: 10.1016/bs.apar.2015.06.001

Klompmaker AA, Portell RW, Frick MG (2017) Comparative experimental taphonomy of eight marine arthropods indicates distinct differences in preservation potential. Palaeontology 60:773–794. doi: 10.1111/pala.12314

Lee IO, Bae HJ, Kim H-G, Oh C-W (2016) Effects of infestation by *Argeia pugettensis* Dana, 1853 (Isopoda, Bopyridae) on the growth and reproduction of the sand shrimp *Crangon hakodatei* Rathbun, 1902 (Decapoda, Crangonidae) in South Korea. Crustaceana 89:685–699. doi: 10.1163/15685403-00003556

Lester RJG (2005) Isopoda (isopods). In: Rohde K (ed) Marine Parasitology. CSIRO Publishing, Victoria, pp 138–144

Mantelatto F, Miranda I (2010) Temporal dynamic of the relationship between the parasitic isopod *Aporobopyrus curtatus* (Crustacea: Isopoda: Bopyridae) and the anomuran crab *Petrolisthes armatus* (Crustacea: Decapoda: Porcellanidae) in southern Brazil. Latin American Journal of Aquatic Research 38:210–217. doi: 10.3856/vol38-issue2-fulltext-5

Markham JC (1986) Evolution and zoogeography of the Isopoda Bopyridae, parasites of Crustacea Decapoda. In: Gore RH, Heck KL (eds) Crustacean Issues 4. Crustacean Biogeography. A.A. Balkema, Rotterdam, pp 143–164

Markham JC (1978) Bopyrid isopods parasitizing hermit crabs in the northwestern Atlantic Ocean. Bulletin of Marine Science 28:102–117

Markham JC, Dworschak PC (2005) A new species of *Entophilus* Richardson, 1903 (Isopoda: Bopyridae: Entophilinae) from the Gulf of Aqaba, Jordan. Journal of Crustacean Biology 25:413–419. doi: 10.1651/C-2566

McDermott JJ (1991) Incidence and host-parasite relationship of *Leidya bimini* (Crustacea, Isopoda, Bopyridae) in the brachyuran crab *Pachygrapsus transversus* from Bermuda. Ophelia 33:71–95

McDermott JJ (2002) Relationships between the parasitic isopods *Stegias clibanarii* Richardson, 1904 and *Bopyrissa wolffi* Markham, 1978 (Bopyridae) and the intertidal hermit crab *Clibanarius tricolor* (Gibbes, 1850) (Anomura) in Bermuda. Ophelia 56:33–42. doi: 10.1080/00785236.2002.10409487

McDermott JJ (2009) Hypersymbioses in the pinnotherid crabs (Decapoda: Brachyura: Pinnotheridae): a review. Journal of Natural History 43:785–805. doi: 10.1080/00222930802702480

McDermott JJ, Williams JD, Boyko CB (2010) The unwanted guests of hermits: a global review of the diversity and natural history of hermit crab parasites. Journal of Experimental Marine Biology and Ecology 394:2–44

McGrew M, Hultgren KM (2011) Bopyrid parasite infestation affects activity levels and morphology of the eusocial snapping shrimp *Synalpheus elizabethae*. Marine Ecology Progress Series 431:195–204. doi: 10.3354/meps09123

Miyashita Y (1941) Observations on an entoniscid parasite of *Eriocheir japonicus* de Haan, *Entionella fluviatilis* n. g., n. sp. Japanese Journal of Zoology 9:251–267

Moericke W (1889) Die Crustaceen der Stramberger Schichten. Palaeontographica, Supplement 2 6:43–72, pl. 6

Mushtaq S, Shafique S, Khatoon Z (2016) *Micippion asymmetricus*: an entoniscid parasite of the coral crab, *Charybdis feriatus* (Linnaeus, 1758). Pakistan Journal of Zoology 48:1607–1611

Nagler C, Høeg JT, Haug C, Haug JT (2017a) A possible 150 million years old cirripede crustacean nauplius and the phenomenon of giant larvae. Contributions to Zoology 86:213–227

Nagler C, Hörnig MK, Haug JT, Noever C, Høeg JT, Glenner H (2017b) The bigger, the better? Volume measurements of parasites and hosts: Parasitic barnacles (Cirripedia, Rhizocephala) and their decapod hosts. PLOS ONE 12:e0179958. doi: 10.1371/journal.pone.0179958

Néraudeau D, Perrichot V, Batten DJ, Boura A, Girard V, Jeanneau L, Nohra YA, Polette F, Martin SS, Saint Martin J-P, Thomas R (2017) Upper Cretaceous amber from Vendée, north-western France: Age dating and geological, chemical, and palaeontological characteristics. Cretaceous Research 70:77–95. doi: 10.1016/j.cretres.2016.10.001

Neves CA, Santos EA, Bainy ACD (2000) Reduced superoxide dismutase activity in *Palaemonetes argentinus* (Decapoda, Palemonidae) infected by *Probopyrus ringueleti* (Isopoda, Bopyridae). Diseases of Aquatic Organisms 39:155–158. doi: 10.3354/dao039155

O’Brien J, Van Wyk P (1985) Effects of crustacean parasitic castrators (epicaridean isopods and rhizocephalan barnacles) on growth of crustacean hosts. In: Wenner AM (ed) Factors in adult growth, Crustacean Issues 3. A.A. Balkema, Rotterdam, pp 191–218

Oliveira E, Masunari S (1998) Population relationships between the parasite *Aporobopyrus curtatus* (Richardson, 1904) (Isopoda: Bopyridae) and one of its porcelain crab hosts *Petrolisthes armatus* (Gibbes, 1850) (Decapoda: Porcellanidae) from Farol Island, southern Brazil. Journal of Natural History 32:1707–1717. doi: 10.1080/00222939800771221

Owens L, Glazebrook JS (1985) The biology of bopyrid isopods parasitic on commercial penaeid prawns in northern Australia. In: Rothlisberg PC, Hill BJ, Staples DJ (eds) Second Australian National Prawn Seminar. NPS2, Cleveland, Queensland, pp 105–113

Patrulius D (1966) Les Décapodes du Tithonique Inférieur de Woźniki (Carpates Polonaises Occidentales). Annales de la Société Géologique de Pologne 36:495–517, pls. 30–31

Petrić M, Ferri J, Mladineo I (2010) Growth and reproduction of *Munida rutllanti* (Decapoda: Anomura: Galatheidae) and impact of parasitism by *Pleurocrypta* sp. (Isopoda: Bopyridae) in the Adriatic Sea. Journal of the Marine Biological Association of the United Kingdom 90:1395–1404. doi: 10.1017/S0025315409991615

Poinar G, Duarte D, Santos MJ (2010) *Halomonhystera parasitica* n. sp. (Nematoda: Monhysteridae), a parasite of *Talorchestia brito* (Crustacea: Talitridae) in Portugal. Systematic Parasitology 75:53–58. doi: 10.1007/s11230-009-9210-x

Portell RW, Collins JSH (2004) Decapod crustaceans of the Lower Miocene Montpelier Formation, White Limestone Group of Jamaica. Cainozoic Research 3:109–126

Poulin R (2011) Evolutionary Ecology of Parasites, 2nd edition. Princeton University Press, Princeton

Poulin R, Krasnov BR, Mouillot D (2011) Host specificity in phylogenetic and geographic space. Trends in Parasitology 27:355–361. doi: 10.1016/j.pt.2011.05.003

Poulin R, Mouillot D (2005) Combining phylogenetic and ecological information into a new index of host specificity. Journal of Parasitology 91:511–514. doi: 10.1645/GE-398R

Radwański A (1972) Isopod-infected prosoponids from the Upper Jurassic of Poland. Acta Geologica Polonica 22:499–506

Rasch JA, Bauer RT (2015) Temporal variation in population structure of the isopod *Urobopyrus processae* Richardson, 1904 (Isopoda: Bopyridae) infesting the branchial chamber of the night shrimp *Ambidexter symmetricus* Manning and Chace, 1971 (Decapoda: Processidae). Nauplius 23:89–103. doi: 10.1590/S0104-64972015002317

Rasmussen E (1973) Systematics and ecology of the Isefjord marine fauna (Denmark) with a survey of the eelgrass *(Zostera)* vegetation and its communities. Ophelia 11:1–507. doi: 10.1080/00785326.1973.10430115

Reinhard EG (1956) Parasitic castration of Crustacea. Experimental Parasitology 5:79–107. doi: 10.1016/0014-4894(56)90007-8

Remeš M (1921) Excroissance des crustacés du Tithonique de Štramberk. Bulletin international de l’Académie des Sciences de Bohême 23:36–37, pl. 1

Reuss AE (1858) Über kurzschwänzige Krebse im Jurakalke Mährens. Sitzungsberichte der Kaiserlichen Akademie der Wissenschaften, (Mathematischnaturwissenschaftliche Classe) 31:5–13

Robins CM, Feldmann RM, Schweitzer CE (2013) Nine new genera and 24 new species of the Munidopsidae (Decapoda: Anomura: Galatheoidea) from the Jurassic Ernstbrunn Limestone of Austria, and notes on fossil munidopsid classification. Annalen des Naturhistorischen Museums in Wien, Serie A 115:167–251

Roccatagliata D, Lovrich GA (1999) Infestation of the false king crab *Paralomis granulosa* (Decapoda: Lithodidae) by *Pseudione tuberculata* (Isopoda: Bopyridae) in the Beagle Channel, Argentina. Journal of Crustacean Biology 19:720–729

Rohde K (1980) Host specificity indices of parasites and their application. Experientia 36:1369–1371. doi: 10.1007/BF01960103

Rohde K, Rohde PP (2005) The ecological niches of parasites. In: Rohde K (ed) Marine Parasitology. CSIRO Publishing, Collingwood, pp 286–293

Rollins HB, Brezinski DK (1988) Reinterpretation of crinoid-platyceratid interaction. Lethaia 21:207–217. doi: 10.1111/j.1502-3931.1988.tb02072.x

Román-Contreras R, Romero-Rodríguez J (2013) Prevalence and reproduction of *Bopyrina abbreviata* (Isopoda, Bopyridae) in Laguna de Términos, SW Gulf of Mexico. Journal of Crustacean Biology 33:641–650. doi: 10.1163/1937240X-00002182

Romero-Rodríguez J, Román-Contreras R (2011) Changes in secondary sexual characters of males of *Thor floridanus* (Decapoda, Hippolytidae), infested by *Bopyrinella thorii* (Isopoda, Bopyridae). Crustaceana 84:1041–1050. doi: 10.1163/001121611X586693

Romero-Rodríguez J, Román-Contreras R (2008) Aspects of the reproduction of *Bopyrinella thorii* (Richardson, 1904) (Isopoda, Bopyridae), a branchial parasite of *Thor floridanus* Kingsley, 1878 (Decapoda, Hippolytidae) in Bahía de la Ascensión, Mexican Caribbean. Crustaceana 81:1201–1210. doi: 10.1163/156854008X374522

Romero-Rodríguez J, Román-Contreras R, Cházaro-Olvera S, Martínez-Muñoz MA (2016) Growth of individuals within the parasite-host association *Bopyrina abbreviata* (Isopoda, Bopyridae) and *Hippolyte zostericola* (Decapoda, Caridea), and variations in parasite morphology. Invertebrate Reproduction & Development 60:39–48. doi: 10.1080/07924259.2015.1126536

Ruiz de Gaona M (1943) Nota sobre Crustáceos Decápodos de la cantera del Monte Orobe (Alsasua). Boletin Real Sociedad Española de Historia Natural 40:425–433, pl. 28

Ruiz GM, Lindberg DR (1989) A fossil record for trematodes: extent and potential uses. Lethaia 22:431–438. doi: 10.1111/j.1502-3931.1989.tb01447.x

Schädel M, Nagler C, Haug JT (2018) Taking a deeper look – a fossil isopod revisited by ‘virtual paleontology.’ In: 9th International Crustacean Congress, Abstract Book. Washington, D. C., p 60

Schneider S, Harzhauser M, Kroh A, Lukeneder A, Zuschin M (2013) Ernstbrunn Limestone and Klentnice beds (Kimmeridgian Berriasian; Waschberg-dánice Unit; NE Austria and SE Czech Republic): state of the art and bibliography. Bulletin of Geosciences 88:105–130

Scholtz G, Ponomarenko E, Wolff C (2009) Cirripede cleavage patterns and the origin of the Rhizocephala (Crustacea: Thecostraca). Arthropod Systematics & Phylogeny 67:219–228

Schweitzer CE, Feldmann RM (2009a) Revision of the genus Cyclothyreus Remeš, 1895 (Decapoda: Brachyura: Dromioidea). Neues Jahrbuch für Geologie und Paläontologie - Abhandlungen 253:357–372

Schweitzer CE, Feldmann RM (2009b) Revision of the Prosopinae sensu Glaessner, 1969 (Crustacea: Decapoda: Brachyura) including four new families, four new genera, and five new species. Annalen des Naturhistorischen Museums in Wien, Serie A 110:55–121

Schweitzer CE, Feldmann RM (2010) Revision of Cycloprosopon and additional notes on Eodromites (Brachyura: Homolodromioidea: Goniodromitidae). Annalen des Naturhistorischen Museums in Wien Serie A 112:169–194

Schweitzer CE, Feldmann RM, Franţescu OD, Klompmaker A (2012) Revision of Etyidae Guinot and Tavares, 2001 (Crustacea: Brachyura). Journal of Paleontology 86:129–155. doi: 10.1666/11-060.1

Seguenza G (1876) Ricerche Paleontologiche intorno ai Cirripedi Terziarii della Provincia di Messina. Atti Dell’Accademia Pontaniana 3:268–481

Serrano-Sánchez M de L, Nagler C, Haug C, Haug JT, Centeno-García E, Vega FJ (2016) The first fossil record of larval stages of parasitic isopods: cryptoniscus larvae preserved in Miocene amber. Neues Jahrbuch für Geologie und Paläontologie - Abhandlungen 279:97–106. doi: 10.1127/njgpa/2016/0543

Sherman MB, Curran MC (2013) The effect of the bopyrid isopod *Probopyrus pandalicola* (Packard, 1879) (Isopoda, Bopyridae) on the survival time of the daggerblade grass shrimp *Palaemonetes pugio* Holthuis, 1949 (Decapoda, Palaemonidae) during starvation at two different temperatures. Crustaceana 86:1328–1342. doi: 10.1163/15685403-00003245

Shields JD, Kuris AM (1985) Ectopic infections of *Portunion conformis* (Isopoda: Entoniscidae) in *Hemigrapsus* spp. Journal of Invertebrate Pathology 45:122–124

Shiino SM (1942) On the parasitic isopods of the family Entoniscidae, especially those found in the vicinity of Seto. Memoirs of the College of Science, Kyoto Imperial University, Series B 17:37–76

Siveter DJ, Briggs DEG, Siveter DJ, Sutton MD (2015) A 425-million-year-old Silurian pentastomid parasitic on ostracods. Current Biology 25:1632–1637. doi: 10.1016/j.cub.2015.04.035

Smit NJ, Bruce NL, Hadfield KA (2014) Global diversity of fish parasitic isopod crustaceans of the family Cymothoidae. International Journal for Parasitology: Parasites and Wildlife 3:188–197. doi: 10.1016/j.ijppaw.2014.03.004

Smith AE, Chapman JW, Dumbauld BR (2008) Population structure and energetics of the bopyrid isopod parasite *Orthione griffenis* in mud shrimp *Upogebia pugettensis*. Journal of Crustacean Biology 28:228–233. doi: 10.1651/0278-0372(2008)028[0228:PSAEOT]2.0.CO;2

Smith JA, Handley JC, Dietl GP (2018) On drilling frequency and Manly’s alpha: towards a null model for predator preference in paleoecology. PALAIOS 33:61–68. doi: 10.2110/palo.2017.098

Somers IF, Kirkwood GP (1991) Population ecology of the grooved tiger prawn, *Penaeus semisulcatus*, in the north-western Gulf of Carpentaria, Australia: growth, movement, age structure and infestation by the bopyrid parasite *Epipenaeon ingens*. Marine and Freshwater Research 42:349–367. doi: 10.1071/MF9910349

Takahashi T, Matsuura S (1994) Laboratory studies on molting and growth of the shore crab, *Hemigrapsus sanguineus* de Haan, parasitized by a rhizocephalan barnacle. The Biological Bulletin 186:300–308. doi: 10.2307/1542276

Tapanila L (2008) Direct evidence of ancient symbiosis using trace fossils. The Paleontological Society Papers 14:271–287. doi: 10.1017/S1089332600001728

Torchin ME, Lafferty KD, Kuris AM (2001) Release from parasites as natural enemies: increased performance of a globally introduced marine crab. Biological Invasions 3:333–345

Trilles J-P, Hipeau-Jacquotte R (2012) Symbiosis and parasitism in the Crustacea. In: Charmantier-Daures Forest (†) M, Vaupel Klein Schram C, Charmantier-Daures Forest (†) M, Vaupel Klein Schram C (eds) Treatise on Zoology - Anatomy, Taxonomy, Biology. The Crustacea, Volume 3. Brill, Leiden, pp 239–317

Upeniece I (2001) The unique fossil assemblage from the Lode Quarry (Upper Devonian, Latvia). Mitteilungen aus dem Museum für Naturkunde in Berlin – Geowissenschaftliche Reihe 4:101–119. doi: 10.1002/mmng.20010040108

Upeniece I (2011) Palaeoecology and juvenile individuals of the Devonian placoderm and acanthodian fishes from Lode site, Latvia. Dissertation, University of Latvia

Van Straelen V (1940) Crustacés Décapodes nouveaux du Crétacique de la Navarre. Bulletin du Musée Royal d’Histoire Naturelle de Belgique 16:1–5, pl. 1

Van Straelen V (1931) Crustacea Eumalacostraca (Crustaceis decapodis exclusis). In: Quenstedt W (ed) Fossilium Catalogus, I Animalia. Part 48. W. Junk, Berlin, pp 1–98

Van Straelen V (1928) Contribution à l’étude des Isopodes Méso- et Cénozoïques. Mémoires Academie Royale des Belgique, Classe de Sciences Collection, Second Series 9:1–68

Van Wyk PM (1982) Inhibition of the growth and reproduction of the porcellanid crab *Pachycheles rudis* by the bopyrid isopod, *Aporobopyrus muguensis*. Parasitology 85:459–473. doi: 10.1017/S0031182000056249

Vannier J, Abe K (1993) Functional morphology and behavior of *Vargula hilgendorfii* (Ostracoda: Myodocopida) from Japan, and discussion of its crustacean ectoparasites: preliminary results from video recordings. Journal of Crustacean Biology 13:51–76. doi: 10.1163/193724093X00444

Vía Boada L (1981) Les Crustacés Décapodes du Cénomanien de Navarra (Espagne): premiers resultats de l’étude des Galatheidae. Géobios 14:247–251

von Meyer H (1835) Briefliche Mittheilungen. Neues Jahrbuch für Mineralogie, Geologie, Geognosie und Petrefaktenkunde 1835:329

von Meyer H (1842) Über die in dem dichten Jurakalk von Aalen in Würtemburg vorkommenden Spezies des Crustaceengenus Prosopon. Beiträge zur Petrefaktenkunde 5:70–75, pl. 15

von Meyer H (1857) Briefliche Mittheilungen. Neues Jahrbuch für Mineralogie, Geognosie, Geologie und Petrefaktenkunde 1857:556

von Meyer H (1858) Briefliche Mittheilungen. Neues Jahrbuch für Mineralogie, Geognosie, Geologie und Petrefaktenkunde 1858:59–62

von Meyer H (1864) Briefliche Mittheilungen. Neues Jahrbuch für Mineralogie, Geognosie, Geologie und Petrefaktenkunde 1864:20

Walker SP (1977) *Probopyrus pandalicola*: Discontinuous ingestion of shrimp hemolymph. Experimental Parasitology 41:198–205. doi: 10.1016/0014-4894(77)90145-X

Wehner G (1988) Über die Prosoponiden (Crustacea, Decapoda) des Jura. Dissertation, Ludwig-Maximilians-Universität München

Weitschat W, Guhl W (1994) Erster Nachweis fossiler Ciliaten: first fossil Ciliates. Paläontologische Zeitschrift 68:17–31. doi: 10.1007/BF02989430

Wenner EL, Windsor NT (1979) Parasitism of galatheid crustaceans from the Norfolk Canyon and Middle Atlantic Bight by bopyrid isopods. Crustaceana 37:293–303. doi: 10.1163/156854079X01176

Wienberg Rasmussen H, Jakobsen SL, Collins JSH (2008) Raninidae infested by parasitic Isopoda (Epicaridea). Bulletin of the Mizunami Fossil Museum 34:31–49

Williams JD, Boyko CB (2012) The global diversity of parasitic isopods associated with crustacean hosts (Isopoda: Bopyroidea and Cryptoniscoidea). PLoS One 7:e35350. doi: 10.1371/journal.pone.0035350

Xylander W (2005) Gyrocotylidea (unsegmented tapeworms). In: Xylander W (ed) Marine Parasitology. CABI Publishing, Oxon

Yamaguti S (1963) Parasitic Copepoda and Branchiura of fishes. Interscience Publishers, New York

Yasuoka N, Yusa Y (2017) Effects of a crustacean parasite and hyperparasite on the Japanese spiny oyster *Saccostrea kegaki*. Marine Biology 164:217. doi: 10.1007/s00227-017-3250-6

Zeiss A (2001) Die Ammonitenfauna der Tithonklippen von Ernstbrunn, Niederösterreich. Neue Denkschriften des Naturhistorischen Museums in Wien, Neue Serie 6:1–117

